# Artificial Intelligence Powered Biomarker Discovery: A Large-Scale Analysis of 236 Studies Across 19 Therapeutic Areas and 147 Diseases

**DOI:** 10.1101/2025.08.28.672795

**Authors:** Ahtisham Fazeel Abbasi, Shiza Naveed, Muhammad Nabeel Asim, Muhammad Sajjad, Sebastian Vollmer, Andreas Dengel

## Abstract

Biomarkers are the molecular signatures that drive and reflect disease states and are indispensable for disease diagnosis, therapeutic target identification, and drug development. The landscape of biomarker discovery has undergone a transformative shift with the emergence of AI-powered predictive pipelines that can integrate complex, high-dimensional datasets. However, the field still lacks a comprehensive, cross-disciplinary foundation that unites AI pipelines with disease-specific biological insights. Together, a combined scattered knowledge of 15 review articles fails to provide a unified framework encompassing data availability, methodological trends, and disease-specific biomarker discoveries across therapeutic areas. Most prior efforts have concentrated on narrow aspects, either focusing on disease-specific AI models or individual stages of the biomarker discovery pipelines, leaving a substantial gap in translational utility. This study addresses this gap by systematically consolidating and analyzing findings from 236 AI-driven biomarker discovery studies. We systematically map the trends of datasets, data modalities, preprocessing strategies, feature engineering methods, AI models, and explainability methods across 147 diseases, which we further organize into 19 therapeutic areas. By doing so, we aim to provide a comprehensive resource that not only highlights current trends and gaps but also lays the groundwork for future advancements, including the design of multi-task learning models and multimodal AI frameworks tailored to complex biomedical data.

## 1 Introduction

Biomarkers are measurable indicators that are used to discern normal and abnormal biological processes such as disease states and therapeutic responses [1]. Typically, they can be detected in saliva [2], blood, urine, tissue, or cerebrospinal fluid and are constituted by DNA, RNA, proteins, or other biomolecules [3–5]. Figure 1 shows their categorization into 7 groups according to their clinical applications [6]: diagnostic (tau protein for Alzheimer’s disease) [7], monitoring (creatinine for renal function) [8], pharmacodynamic (ctDNA in cancer treatment), predictive (HER2 in breast cancer), prognostic (CAG repeats in Huntington’s disease) [9], safety (Serum creatinine levels in kidney toxcity), and susceptibility (IL-6 in schizophrenia) [10].

**Fig. 1.**
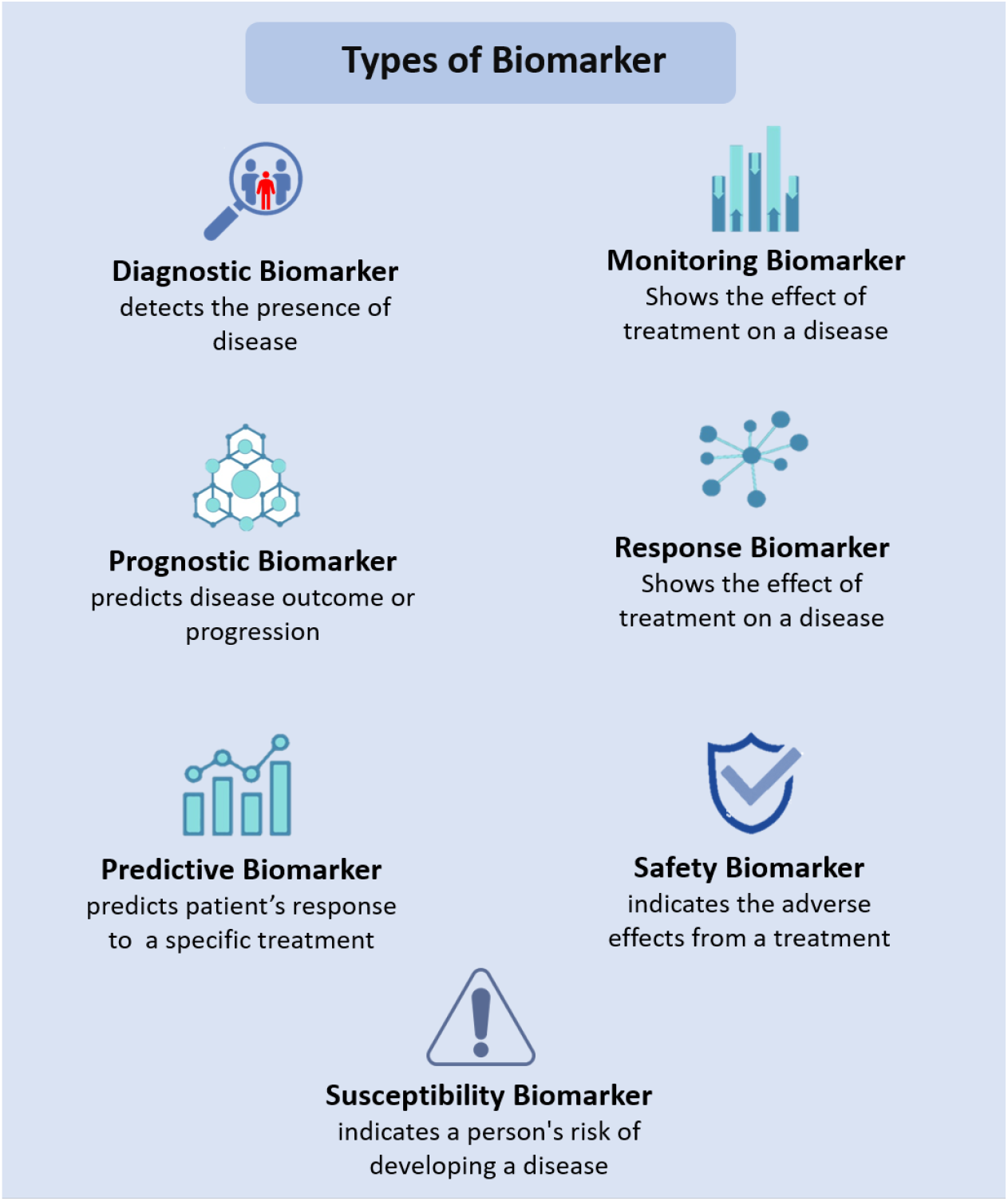
Categorization of biomarkers into 7 groups based on their clinical applications.

Biomarkers are indispensable for disease detection, prognosis, and treatment evaluation, thereby serving a critical function in both clinical practice and research [11–13]. They provide crucial insights into disease mechanisms, facilitating early diagnosis and timely intervention. In precision medicine, biomarkers help tailor treatments to individual biological profiles, improving therapeutic precision [14]. They also support drug development by identifying suitable patients and tracking adverse effects of treatments [15, 16].

Traditional biomarker discovery methods [17– 23] have laid the foundation for clinical research, but face limitations in efficiency, scalability, and integrating complex biological data [24]. These processes often require extensive experimental validation, making it time-consuming and costly [25, 26]. In addition, the identification of biomarkers is further complicated by disease heterogeneity and the interplay of genetic and environmental factors [27].

Building on the success of artificial intelligence (AI) in natural language processing (NLP) and computer vision (CV), researchers are now exploring AI-driven approaches to enhance biomarker discovery [28–30]. With the goal of accelerating and improving the identification of novel biomarkers, AI is being applied to analyze complex biological data [31], integrate multi-omics information [32], and increase predictive accuracy in clinical applications as shown in Figure 2. Particularly, AI is used in biomarker discovery for diverse problem settings, including but not limited to: (a) disease vs healthy classification [33–35], (b) disease molecular subtype classification [36], (c) drug response prediction, and (d) survival analyses.

**Fig. 2.**
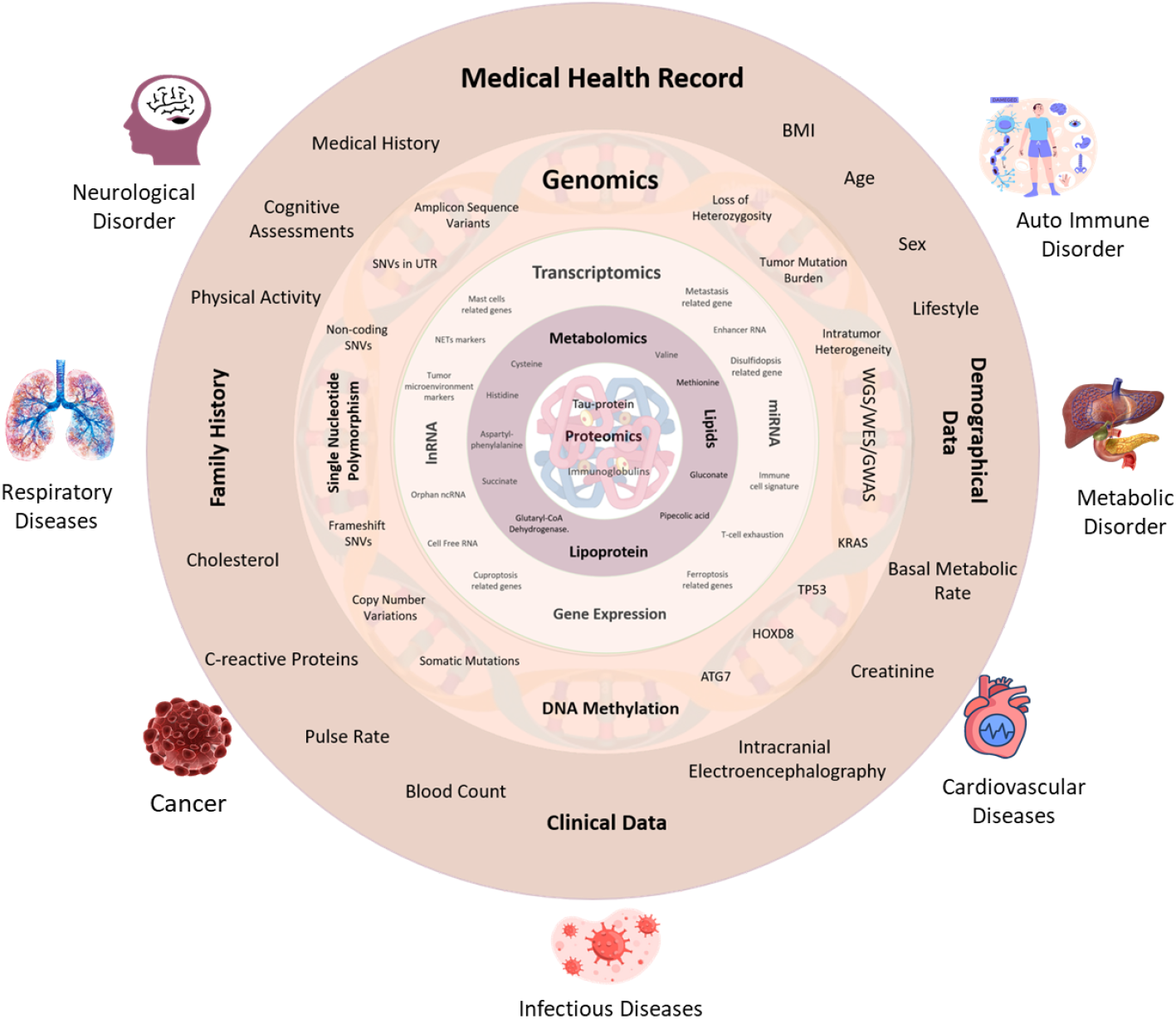
The integration of multimodal data begins with external-level information such as clinical and demographic data, progressively incorporating deeper layers of molecular data—including genomic, transcriptomic, mutation, proteomic, and methylation modalities—to enable comprehensive understanding and precise biomarker discovery.

**Fig. 3.**
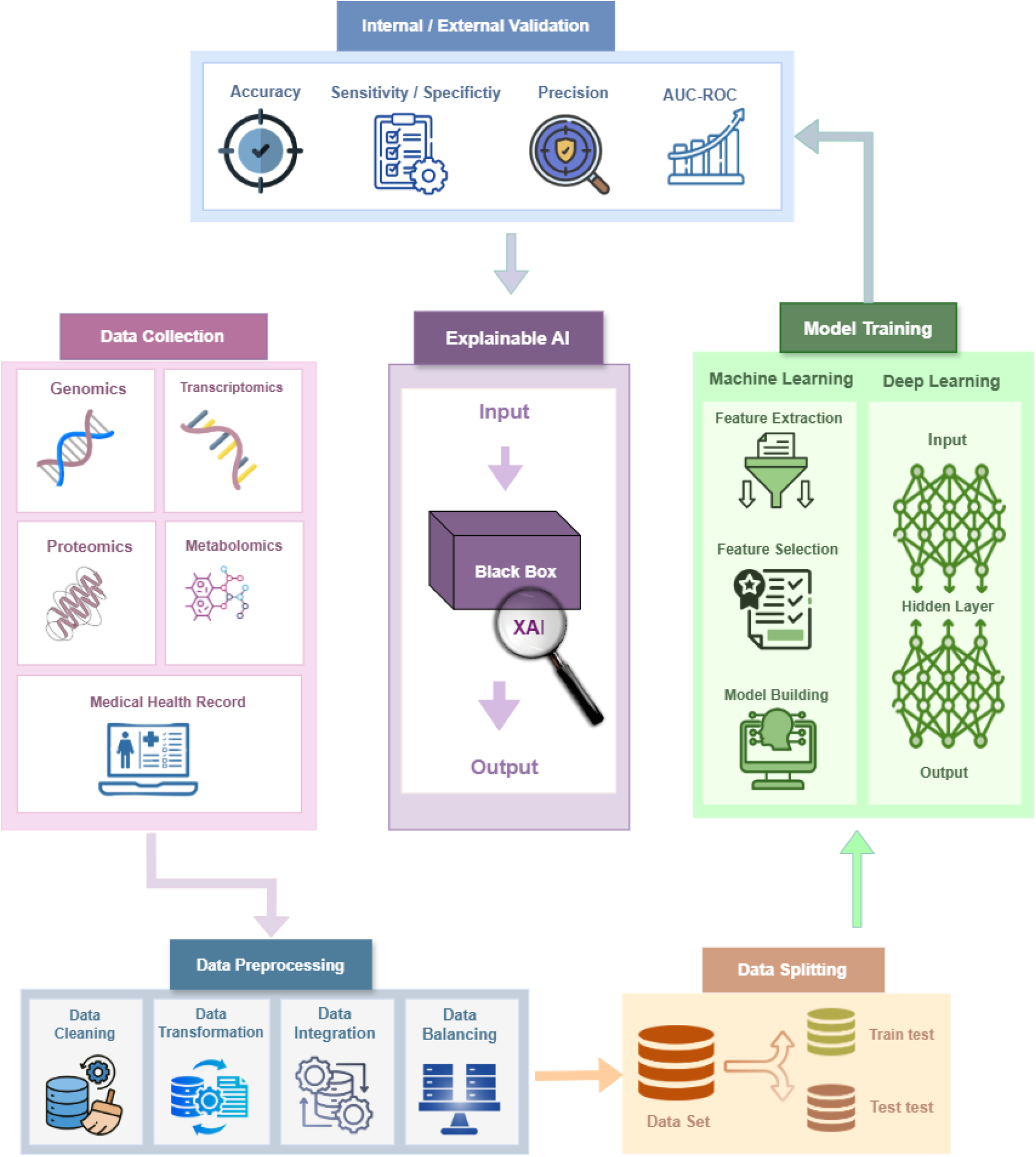
General steps for developing an AI-driven biomarker discovery pipeline.

**Fig. 4.**
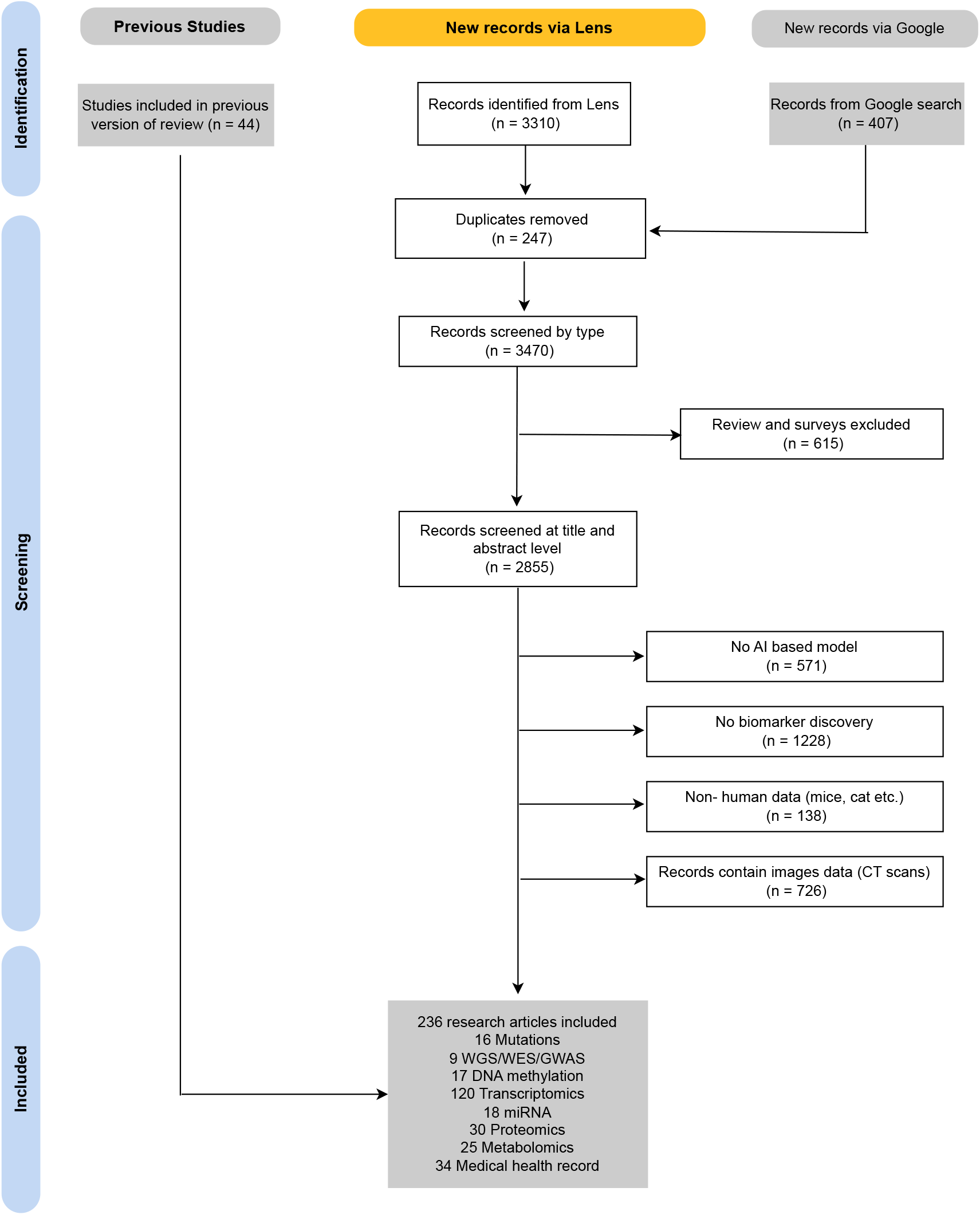
PRISMA flow diagram for study

To accelerate and expedite the development of AI-driven applications for biomarker discovery, apart from the development of 239 task-specific applications, in the last 5 years, 15 review articles have been published. The primary focus of these articles is to summarise trends in problem settings, datasets, data preprocessing and feature engineering methods, data splits, ML/DL models, and XAI in biomarker discovery. However, the focus of these studies is often limited to a single aspect of biomarker discovery, failing to provide a comprehensive overview of biomarker discovery and its relationship to evolving AI trends. While existing reviews are valuable for specific applications, these reviews may not fully explore the potential of AI across the entire biomarker discovery pipeline. More details about existing review papers are presented in Table 1. Following the need for a comprehensive review article for biomarker discovery the contributions of this paper are manifold:

**Table 1.**
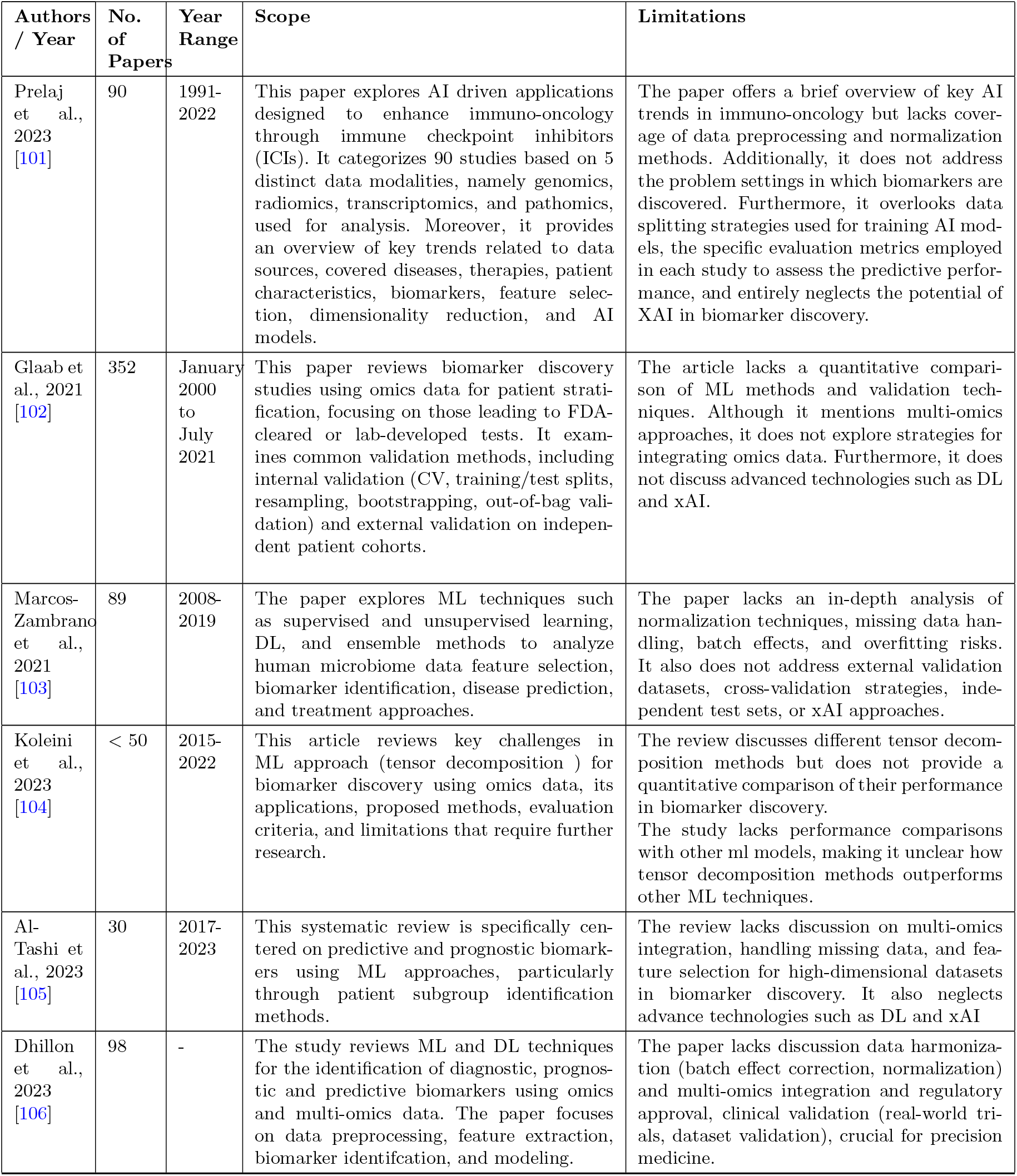
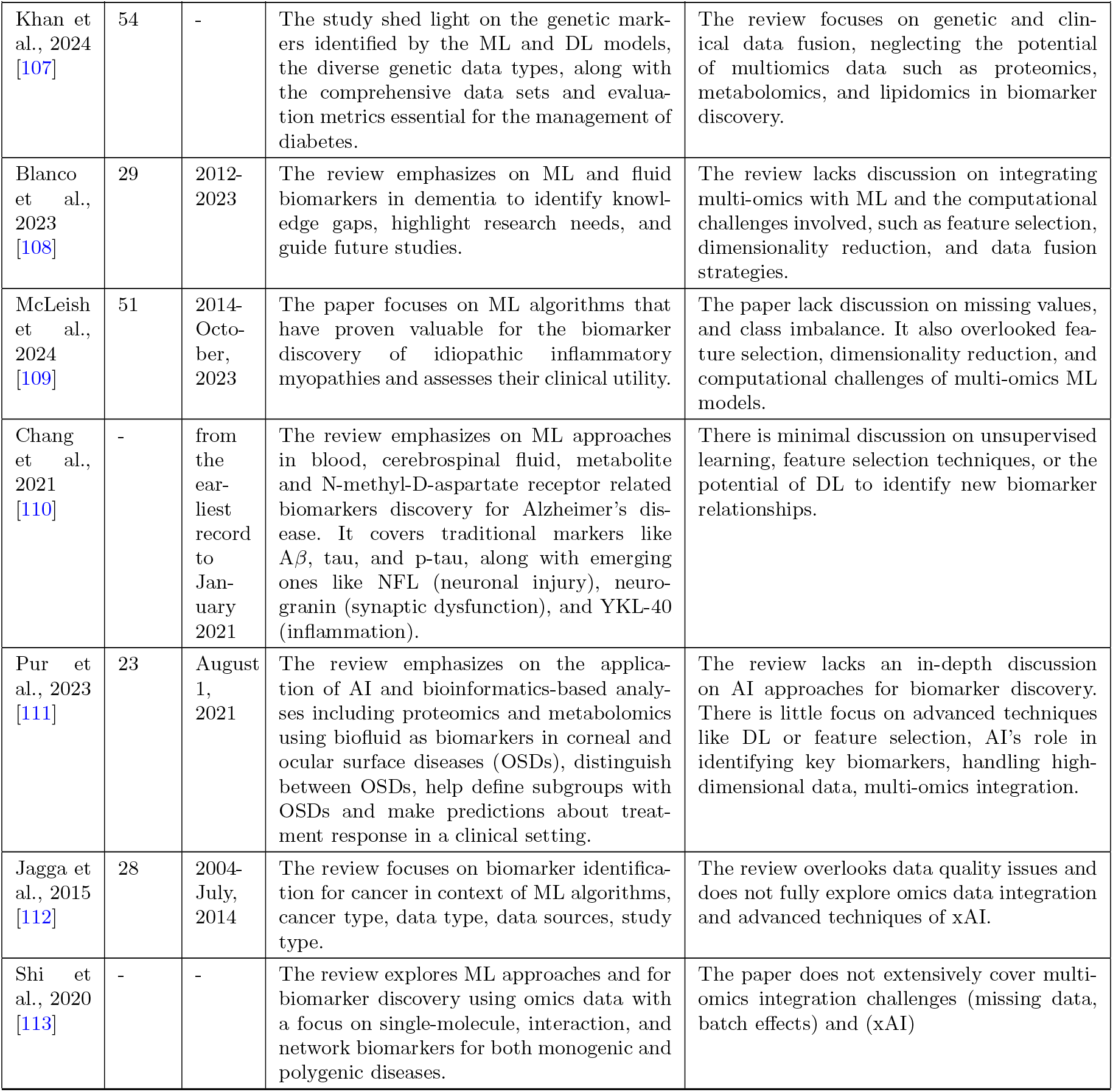
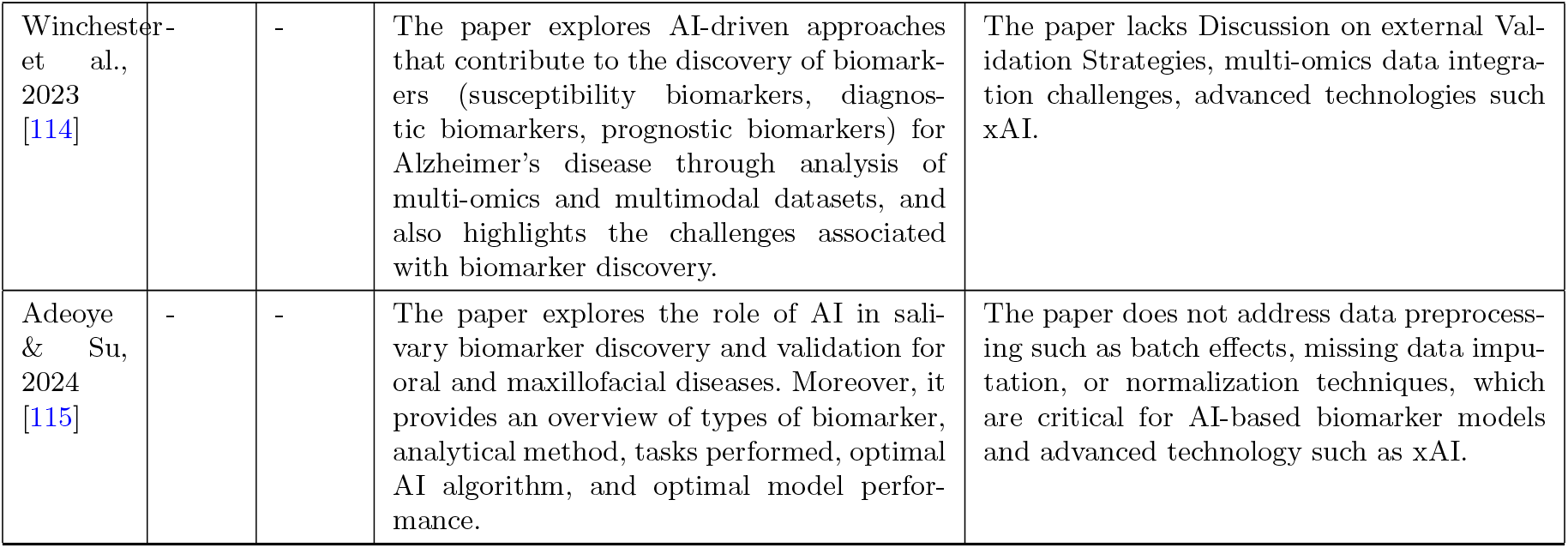
Overview of Scope and Limitations in Existing Survey Papers.

- It brings together 14 diverse review papers related to biomarker discovery and consolidates their analytical frameworks to systematically map research scopes and methodological constraints. The integrated analysis offers researchers a comprehensive reference for navigating current advancement and facilitates efficient access to critical findings and key knowledge gaps in the domain.
- It provides comprehensive details of 236 AI-driven biomarker discovery articles published between 2020 and 2025.

This study systematically delves into diverse aspects of biomarker discovery with 11 diverse research questions (RQs) and objectives: (i) How diverse is the disease coverage in existing research focused on AI-based biomarker discovery? (ii) How do studies address biomarker discovery, ranging from broad examinations of multiple cancer subtypes to focused analyses of individual subtypes? (iii) What are the main problem settings in biomarker discovery, and how are studies allocated across these settings? (iv) Which public and private data sources are most often utilized in biomarker discovery research, what data types do they encompass, and how have studies leveraged these databases? (v) What are the most prevalent data modalities in biomarker discovery, and how are they associated with different diseases? (vi) What is the distribution of studies across various data modalities in biomarker discovery? How have data splitting strategies been used in AI-driven biomarker discovery? (vii) How have feature engineering methods been utilized in biomarker discovery across diverse data modalities and diseases? (viii) Which statistical, machine learning (ML), and deep learning (DL) models are utilized in biomarker discovery for various diseases and problem settings? (ix) Which biomarker discovery studies have openly provided their source code, and what kind of methods are available in open-source prediction tools? (x) What is the clinical and translational impact of biomarkers identified in existing research? (xi) In which journals, omics omics-based biomarker discovery studies have been published?

## 2 Background

AI has transformed biomarker discovery by leveraging high-throughput technologies to decode complex biological datasets [37–39]. Typically, AI-driven biomarker discovery follows a structured pipeline comprising several key stages: data collection, preprocessing, feature engineering, model development, performance evaluation, explainability analysis, and biological interpretation and validation [40]. This section provides technical details of the fundamental components of an AI-driven biomarker discovery pipeline, which serves as a conceptual and practical framework for understanding how AI is applied to identify reliable and interpretable biomarkers in biomedical research.

The first step in biomarker discovery is data collection, where biological samples are obtained from various sources such as blood [41–43], plasma [44–46], serum [47], urine [48, 49], cerebrospinal fluid [50], saliva [51, 52], and tissue biopsies [53]. Afterwards, these samples undergo a series of wet lab experiments, such as nucleic acid extraction, sequencing, mass spectrometry, or methylation profiling, to generate high-dimensional numerical datasets that serve as input for computational analyses. In the subsequent step, genomics, transcriptomics, proteomics, and metabolomics based datasets are stored in public repositories such as The Cancer Genome Atlas (TCGA) [54], Gene Expression Omnibus (GEO) [55], European Genome-phenome Archive (EGA) [56], Proteomics Identifications Database (PRIDE) [57], and Metabolomics Workbench [58].

Raw biological datasets often contain inconsistencies and noise due to technical variability, making preprocessing essential for the development of effective AI-driven applications. Batch effects—arising from differences in equipment, personnel, reagents, or environmental conditions—can obscure true biological signals and lead to misleading conclusions [59–62]. Common correction methods include quantile normalization [63, 64], ComBat (empirical Bayes) [65], and regression-based approaches [66]. Missing values, which frequently occur in high-throughput experiments, must be addressed through data imputation techniques such as k-nearest neighbors (KNN), mean/median substitution, or matrix factorization, to ensure model robustness and prevent information loss. Given the diverse numerical ranges of omics data, normalization and scaling are vital to ensure equal feature contribution during model training. Techniques include minmax scaling [67], Z-score normalization [68], and log transformation for reducing the impact of extreme values, especially in RNA-seq and proteomics datasets [69, 70]. Quantile normalization is particularly effective in harmonizing transcriptomic data distributions [64]. Additionally, clinical and categorical variables, such as mutation status, must be encoded numerically using one-hot, label, or frequency encoding methods [71–74].

Multi-omics datasets often include thousands to millions of features and lower number of samples which can lead to the risk of overfitting due to the curse of dimensionality. In order to manage this “curse of dimensionality” and extract meaningful biological insights, two key strategies are used: dimensionality reduction and feature selection. Dimensionality reduction transforms highdimensional data into a lower-dimensional space while preserving essential information [75]. Common methods include principal component analysis (PCA), which creates orthogonal components from the original features [76], and Autoencoders, which use DL to extract nonlinear patterns [77, 78]. Visualization tools like -distributed stochastic neighbor embedding (t-SNE) and uniform manifold approximation and projection (UMAP) project data into 2D or 3D spaces to reveal hidden structure and clusters [79]. In contrast, feature selection retains the original variables but selects only the most relevant ones for model performance [80, 81]. Methods like recursive feature elimination (RFE) [82], LASSO regression with L1 regularization [83], and random forest feature importance (RFFI) [84] help reduce noise, improve model interpretability, and lower computational cost—while maintaining biological relevance.

To ensure robust model performance, datasets must be appropriately split into training, validation, and test sets. The training set is used for model learning, while the validation set helps tune hyperparameters and prevent overfitting. The test set evaluates the performance of the final model on unseen data [85, 86]. Additionally, cross-validation (e.g., k-fold CV) ensures model generalizability by testing on multiple subsets [87].

A variety of ML and DL models are utilized depending on the problem setting. For instance, in disease subtype classification, supervised learning models such as support vector machine (SVM) [88], decision trees (DT) [89], random forest (RF) [90], and gradient boosting (GB) [91] are commonly used for the identification of molecular subtypes. Similarly, survival analysis spans methods such as Cox Proportional Hazards models and DL-based survival models, to predict prognosis or survival of a patient [92].

Model evaluation ensures biomarker reliability by assessing performance using appropriate evaluation metrics. For classification tasks, accuracy, precision, recall, F1-score, and the area under the receiver operating characteristic curve (ROC-AUC) are commonly used. For imbalanced datasets, the precision-recall (PR) curve provides a more informative evaluation [93–95]. In regression-based biomarker discovery, metrics such as Mean Squared Error (MSE), Mean Absolute Error (MAE), and R-squared (R^2^) assess predictive performance [96]. Survival prediction models rely on concordance index (C-index) and integrated Brier score to evaluate risk prediction accuracy [97, 98].

XAI methods such as shapely additive explanations (SHAP) [99] and Local Interpretable Model-agnostic Explanations (LIME) [100] help interpret feature importance and improve the transparency of biomarker discovery models. XAI aims to provide insights into model decisionmaking, improving transparency and trust. Broadly, explainability techniques fall into posthoc interpretability (e.g., feature attribution, surrogate models) and inherently interpretable models (e.g., DT, generalized linear models). By integrating XAI, AI-driven biomarker discovery becomes more reliable and interpretable for clinical applications.

## 3 Literature Review

In recent years, multiple review papers have been published, with the objective of each review centered on summarizing fundamental concepts in AI-driven biomarker discovery and identifying trends in AI approaches that have been utilized for predicting disease diagnosis, prognosis, and treatment response. Table 1 provides a high-level overview of 14 existing review articles with a focus on their scope and limitations. This comprehensive summary is designed to help researchers efficiently find relevant information in the reviewed articles.

Existing review articles primarily focus on studies related to specific diseases, rather than on a broad spectrum of biomarker discovery, which may limit the generalizability of findings across different conditions. For instance, Prelaj et al. [101] and Jagga et al. [112] conducted an extensive study on biomarker discovery in limited cancers. Khan et al. [107] extended the scope of biomarker discovery to diabetes, McLeish et al. [109] investigated idiopathic inflammatory myopathies, Chang et al. [110] provided valuable insights for Alzheimer’s disease, and Pur et al. [111] focused only on corneal and ocular surface diseases.

Existing review articles also have notable limitations in data pre-processing, feature selection, and dimensionality reduction. Critical challenges such as normalization, data imputation, batch effect correction, and overfitting risks have been overlooked [101, 103, 105, 106, 109, 112]. Furthermore, these studies also lack discussion on external validation datasets, cross-validation strategies, or independent test sets [101, 103, 106, 112].

Beyond data pre-processing challenges, the reviewed studies also exhibit methodological and scope-related limitations. Glaab et al. [102] focused solely on omics-based biomarkers that have received clinical approval, either as laboratory-developed tests or FDA-approved diagnostics, limiting the exploration of emerging molecular signatures. Marcos-Zambrano et al. [103] specifically analyzed human microbiome data, which does not encompass broader omics modalities. Koleini et al. [104] explored tensor decomposition methodologies, narrowing the scope of computational techniques applied to biomarker discovery. Additionally, Al-Tashi et al. [105] centered on recognizing predictive and prognostic biomarkers, whereas Dhillon et al. [106] adopted a more comprehensive approach by addressing diagnostic, prognostic, and predictive biomarker identification. To the best of our knowledge, no existing review or survey provides an indepth analysis of explainability of ML/DL models while highlighting their applications and their importance for biomarker discovery [104, 106, 111, 116]. Few studies, such as [117] highlighted the significance of XAI in biomarker discovery and urge for more research in this domain. Reviews that specifically target XAI applications in biomarker discovery are rare and typically confined to narrow subfields.

The aforementioned limitations of existing review papers underscore the pressing need for a more integrative and standardized review framework for biomarker discovery that systematically encompasses heterogeneous data sources (including multi-omics and clinical data), robust preprocessing strategies, advanced dimensionality reduction and feature selection techniques, a broad spectrum of ML and DL models, and XAI approaches to enhance interpretability, clinical applicability, and translational potential of discovered biomarkers.

## 4 Methodology

The systematic review is conducted following the Preferred Reporting Items for Systematic Reviews and Meta-Analyses (PRISMA) guidelines for structured selection. Relevant articles on biomarker discovery using ML were extracted based on predefined eligibility criteria.

### 4.1 Screening Criteria

A rigorous literature search is conducted using specific keywords in various combinations. The keywords are combined using the Boolean operators such as “AND” and “OR”. Keywords are categorized into 3 categories: (1) Biomarkers—biomarker discovery, biomarker identification, biomarker validation; (2) Omics—genomics, proteomics, transcriptomics, metabolomics; (3) Methodology—AI, ML, and DL. These queries are submitted to literature search engines, such as Lens (https://www.lens.org/) and Google Scholar (https://scholar.google.com/) to retrieve research articles published between January 2020 and December 2024. A total of 3,717 research articles are identified through these queries and then subjected to further screening.

### 4.2 Data selection

To ensure the inclusion of relevant articles, further screening is performed. Following the removal of duplicate studies, a title-based screening is performed based on inclusion/exclusion criteria.

- Studies that focus on ML, DL, and XAI are included.
- Studies excluded that focus on non-human data
- Studies that use images (radiomics, histology, scans) are excluded
- Conference abstracts are excluded.

Following the removal of duplicate studies, 3,490 articles are retained. Title-based selection further narrows the selection to 1,273 articles. Abstract-based screening reduces the count to 352 articles. After full text screening, 236 articles are finalized for inclusion.

## 5 Results

### 5.1 RQ I, II, III: Distribution of Biomarker Discovery Tools Across Diseases and Problem Settings

The primary aim of this section is to present an overview of the distribution of 239 studies/tools across diseases and problem settings. This analysis offers valuable insights into prevailing research trends by highlighting specific diseases that have garnered significant attention. This consolidated overview also serves as a useful resource for researchers seeking to explore biomarker discovery research within their disease of interest. Likewise, distribution analysis of studies across different problem settings enhances our understanding of the current research landscape and also aids in determining underexplored diseases and problem settings, ultimately guiding future research towards high-impact opportunities.

Supplementary Table 1 presents the coverage of diseases across 239 research articles. Particularly, in the past 5 years, 196 tools have been developed to explore a wide range of cancers. Among these, breast cancer (21), lung cancer (35), colorectal cancer (11), gastric/stomach-related cancers (14), and liver cancer types (13) [118– 127]. Other significant cancers include melanoma, gliomas, leukemia, and prostate cancer [128–134]. In contrast, 8 biomarker discovery tools have been developed for pancancer [135–137]. Unlike cancer-specific approaches, pancancer biomarker discovery aims to uncover molecular signatures that are shared across multiple cancers. This strategy leverages data from a broad spectrum of cancers, enabling the identification of common biological pathways and regulatory mechanisms that may not be evident when analyzing each cancer in isolation.

Beyond oncology, biomarker discovery has expanded into numerous non-cancer disease areas. In neurological and mental health disorders, 25 tools have been developed, with Alzheimer’s disease (12) as the major focus. Other conditions like autism, bipolar disorder, and schizophrenia have also gained attention [138–142]. Cardiovascular diseases account for 9 tools, targeting heart failure, myocardial infarction, and aortic dissection [143–145]. Infectious diseases, driven by COVID-19 and tuberculosis, have propelled 15 tools [146–149]. Autoimmune and inflammatory disorders, including ulcerative colitis and rheumatoid arthritis, have led to 10 tools [150–152]. Four tools have addressed reproductive and gynecological diseases like endometriosis and pregnancy loss [134, 153, 154]. Finally, metabolic disorders (6) and hematologic or immunologic conditions (4) such as lymphoma and pheochromocytoma have also been explored [132, 155–159].

Two distinct trends in disease coverage are observed among the reviewed studies. Specifically, 200 studies focus on a single disease, such as a specific cancer or another condition. While many studies are disease-specific, Figure 6 highlights 37 studies that investigate multiple diseases. For instance, Mahdi-Esferizi et al. [134] and Wu et al. [127] each address 13 different cancers, including bladder, breast, and liver cancers. Similarly, Yu et al. [158] explore 12 diseases, encompassing cancers like diffuse large B-cell lymphoma as well as non-cancer conditions such as stroke. Liu et al. (2022) [160] examine 8 mental disorders, including ADHD and depression, while Xie et al. (2024) [132] cover 7 cancer types, such as leukemia and prostate cancer. Additionally, Shomorony et al. (2020) [161] investigate 6 conditions, including diabetes and cardiovascular disease, which highlights the trend of multi-disease studies in advancing cross-disease biomarker insights.

Figure 5 shows the distribution of articles across four distinct problem settings: disease vs. health or subtype classification, survival prediction, disease progression prediction, treatment response prediction, and classification combined with survival prediction. A closer look at Figure 5 reveals that more than 150 studies are centered around disease vs healthy classification or disease subtype classification [148, 150, 162–164]. In addition, around 40 studies chose survival prediction as the main problem setting [119, 130, 165], whereas more than 20 studies also included classification (disease vs healthy, disease subtype classification) [166–168].

**Fig. 5.**
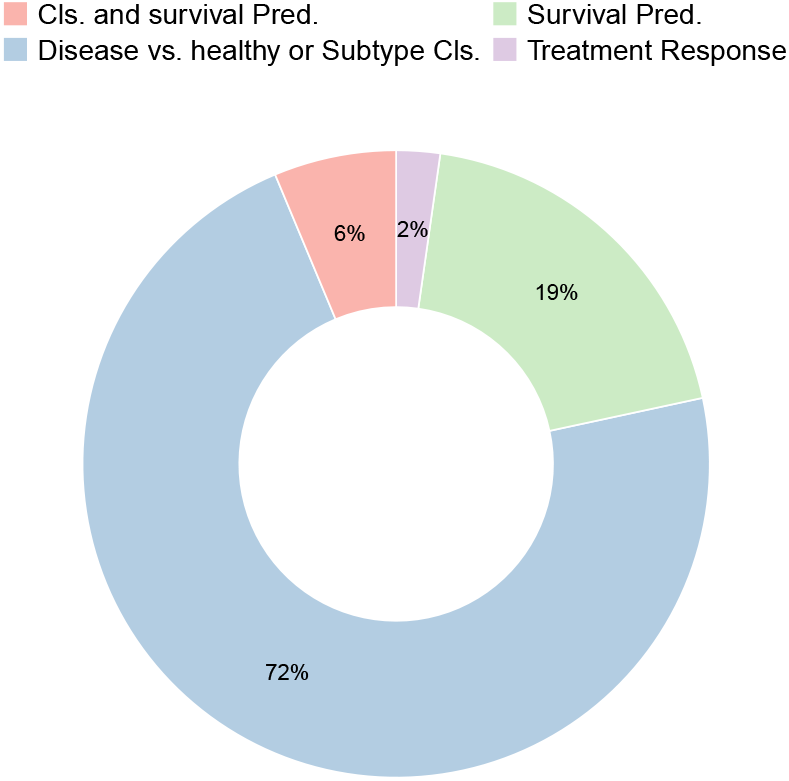
Distribution of studies/tools w.r.t problem settings.

**Fig. 6.**
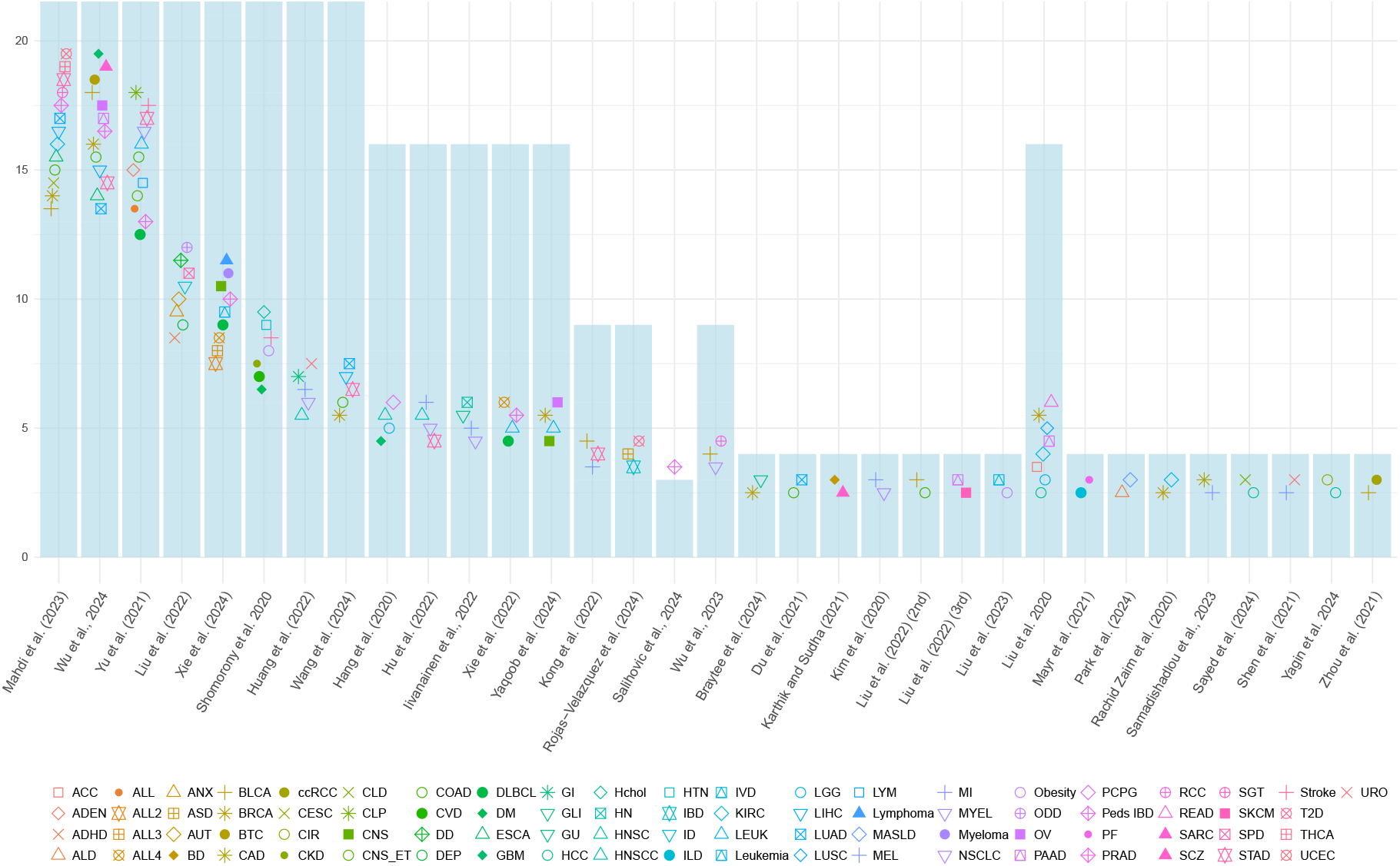
Biomarker discovery studies covering multiple diseases.

**Fig. 7.**
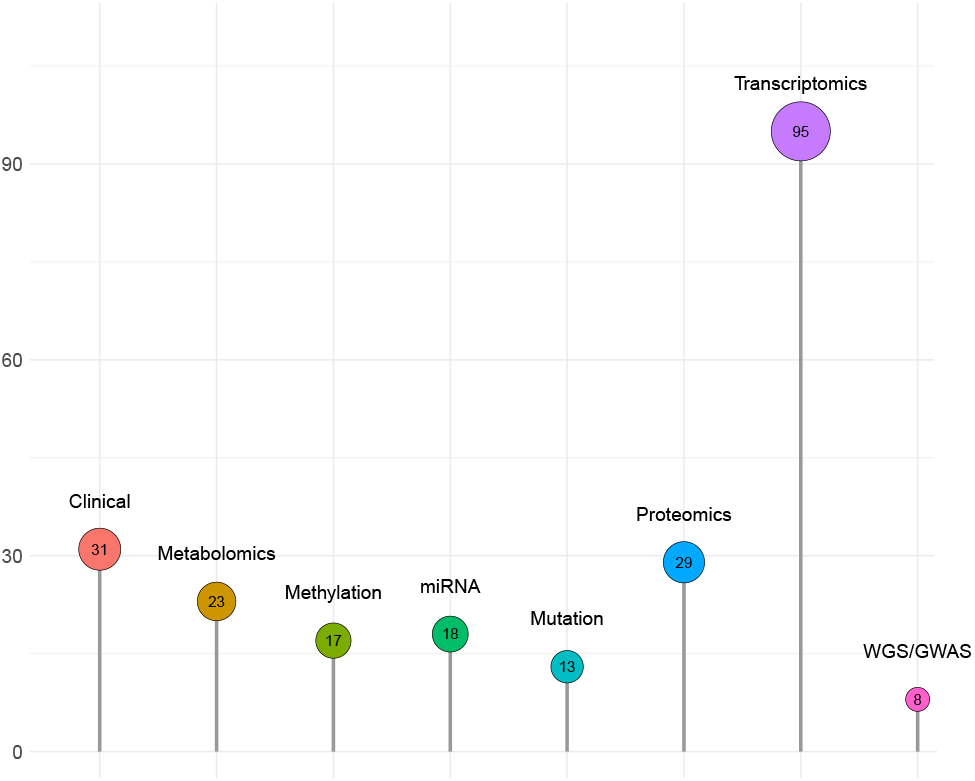
Modality-wise counts of AI-driven biomarker discovery tools/studies.

### 5.2 RQ IV, Biomarker Discovery Data Availability in Public and Private Sources

The development of effective biomarker discovery tools is heavily dependent on the reliability of datasets derived from rigorous wet lab experiments or clinics. These datasets are curated and stored across a range of public repositories, where each dataset entails unique biological and clinical insights. While resources like TCGA [187] specialize in comprehensive cancer-related multi-omics profiles, platforms such as GEO [188] span a wider spectrum of diseases and tissue types. The specialized focus of each database on specific diseases or data modalities necessitates the importance of clearly mapping which resources align with particular research needs. To streamline biomarker discovery tools development efforts, Table 2 provides a curated summary of widely used databases in biomarker discovery, along with details such as types of diseases and omics data covered.

**Table 2.**
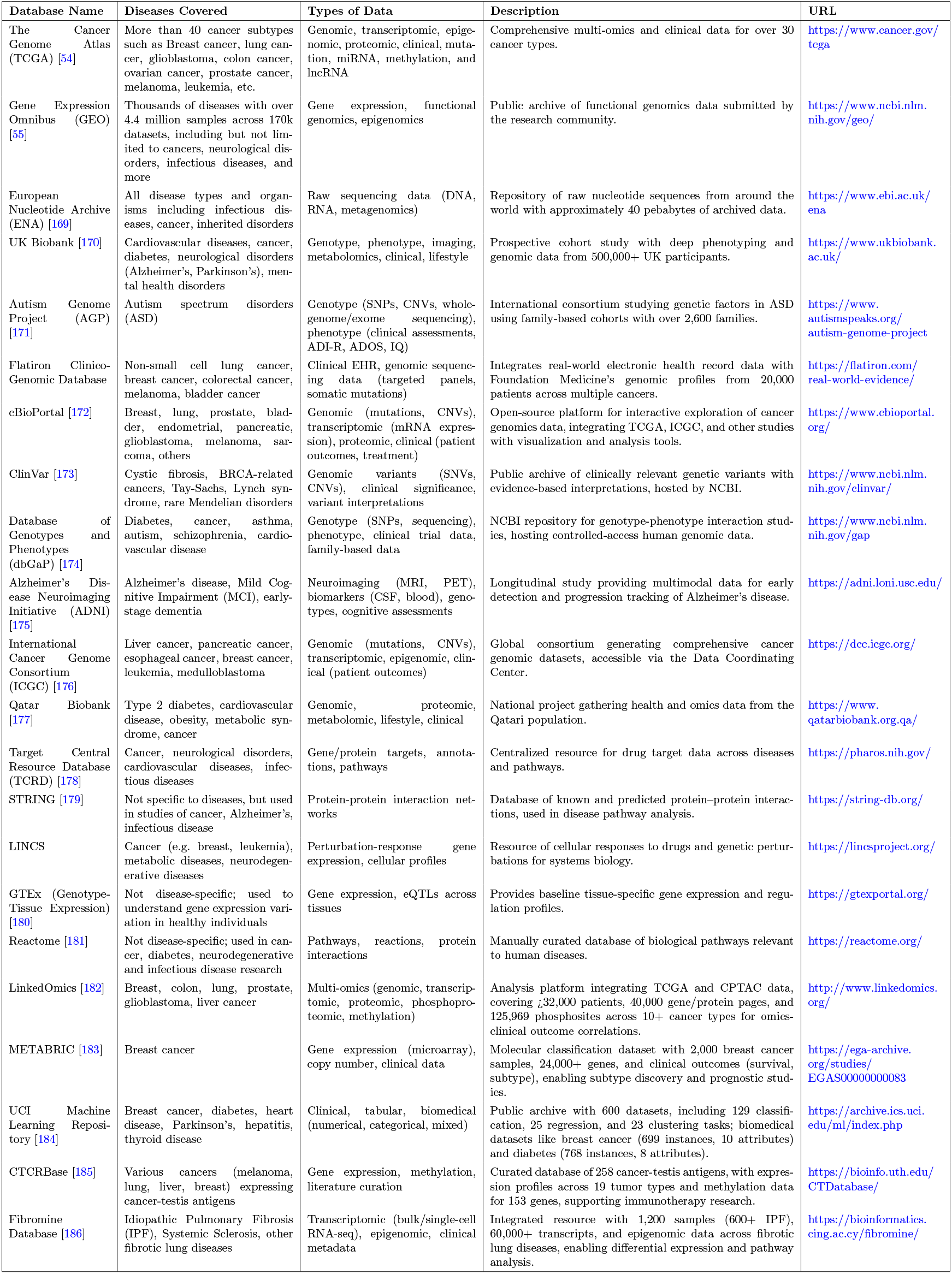
Details of databases used to collect data for biomarker discovery tools.

Supplementary Table 2 presents the utilization of public and private data sources for AI-driven biomarker discovery. Among 236 selected studies, a vast majority opted for publicly available datasets, which are available in public databases such as TCGA and GEO [118, 146, 189–191]. Apart from public databases, a substantial number of biomarker discovery studies have relied on private or institution-specific clinical/omics cohorts. These include data from specialized hospitals and research institutions such as Zhongda Hospital [192], the National Prion Clinic in London [193], Dana-Farber Cancer Institute [194], the Children’s Hospital of Philadelphia (CHOP) [160], and the University of California, San Francisco (UCSF) [195, 196]. Numerous studies also obtained data through ethical approvals from

local hospitals or national health centers, such as TMUGH [197], University of Hong Kong-Shenzhen Hospital [198], and National Taiwan University Hospital [199]. Others utilized biobank-driven or region-specific cohorts like Indivumed (Germany) [200, 201], the Young Finns Study [202], Lupus BioBank of the Upper Rhine Valley [195], and Rheuma-Vor App datasets [195].

The use of publicly available datasets from public databases, namely, TCGA and GEO, offers multiple advantages such as access to large and standardized datasets, enabling reproducibility, cross-study comparisons, and applicability across diverse populations. Conversely, private cohorts from institutions like Zhongda Hospital or biobanks like Indivumed provide tailored, detailed clinical and omics data, as seen in studies like [192] and [200], which can address niche research needs. Yet, they are disadvantaged by smaller sample sizes, restricted access due to ethical or institutional constraints, and potential biases from specific populations, reducing broader applicability.

### 5.3 RQ V, VI: Data Modalities, Their Distribution and Association with Tools and Diseases, and Data Splitting

In pursuit of addressing RQs V and VI, this subsection elucidates prevailing trends in the utilization of data modalities within biomarker discovery. Specifically, for RQ V, it analyzes the distribution of data modalities across 236 studies and explores their tailored usage for specific diseases. In alignment with RQ VI, it also provides a comprehensive analysis of the use of individual data modalities and their combinations, which offers insights and actionable guidance to optimize data selection and integration for enhanced biomarker discovery.

Out of the 236 studies selected for analysis, 31 relied exclusively on clinical data, while the remaining 205 incorporated multiomics data. Multiomics data, which originates from molecular characteristics, is systematically classified into nine distinct categories: gene expression (transcriptomics), microRNA (miRNA), DNA methylation, copy number variation (CNV), whole exome sequencing (WES), whole genome sequencing (WGS), genome-wide association studies (GWAS), metabolomics, and proteomics. Among the 205 studies utilizing multiomics data, 95 utilized transcriptomics data, 29 incorporated proteomics, 23 utilized metabolomics, 13 focused on mutation-based data (CNV) [131, 164, 203, 204], and 35 integrated either DNA methylation [192, 205–207] or miRNA [137, 208–210]. In addition, only a small subset of 8 studies leveraged GWAS, WES, or WGS data [195, 211–213, 213]. Moreover, only 15 studies explored the combined potential of different data modalities, i.e., (methylation, miRNA, transcriptomics, and mutation) [131], and (proteomics, metabolomics, clinical, and WES/WGS/GWAS) 161, which are presented in Table 3.

**Table 3.** Studies that incorporate multiple data modalities for biomarker discovery.

| Modality | [131] | [202] | [195] | [235] | [212] | [236] | [237] | [213] | [196] | [234] | [164] | [238] | [204] | [224] | [239] |
| --- | --- | --- | --- | --- | --- | --- | --- | --- | --- | --- | --- | --- | --- | --- | --- |
| Methylation | ✓ | ✓ | × | × | × | × | ✓ | ✓ | × | ✓ | × | × | × | × | × |
| Clinical | × | ✓ | ✓ | ✓ | ✓ | × | ✓ | ✓ | × | × | ✓ | ✓ | × | × | × |
| Metabolomics | × | ✓ | ✓ | × | × | ✓ | × | × | ✓ | × | × | ✓ | × | × | ✓ |
| miRNA | ✓ | × | × | ✓ | × | × | × | × | × | ✓ | × | × | × | × | × |
| Proteomics | × | × | ✓ | × | × | × | × | × | × | × | × | × | × | × | ✓ |
| Transcriptomics | ✓ | ✓ | × | ✓ | ✓ | ✓ | ✓ | × | ✓ | × | × | × | ✓ | ✓ | × |
| Mutation | ✓ | × | × | × | × | × | ✓ | × | × | × | ✓ | × | ✓ | × | × |
| WGS/WES/GWAS | × | × | ✓ | × | ✓ | × | × | ✓ | × | × | × | × | × | ✓ | × |

The utilization of multiomics data modalities is predominantly observed in cancer research, as evidenced by the disease-wise distribution of omics modalities in Figure 8. This trend reflects the complex molecular landscape of cancers, which necessitates comprehensive profiling to identify robust biomarkers for diagnosis, prognosis, and therapeutic response. Among the various omics modalities, transcriptomics, proteomics, miRNA, and clinical data have been commonly utilized for cancer biomarker discovery. For instance, breast cancer studies leveraged transcriptomics [119, 121, 214], proteomics [215–217], miRNA [127, 210, 218], mutation [131] and clinical data [219, 220]. Similarly, in Non-small cell lung cancer (NSCLC) studies utilized transcriptomics [163, 201, 221], mutation [194] and clinical data [221–223]. In addition, Colorectal cancer research frequently used transcriptomics [121, 123, 224], proteomics [123, 217, 225], and methylation [226]. Moreover, pancancer research also further highlights multi-omics utility i.e., transcriptomics [227, 228], miRNA [137], and mutation data [229].

**Fig. 8.**
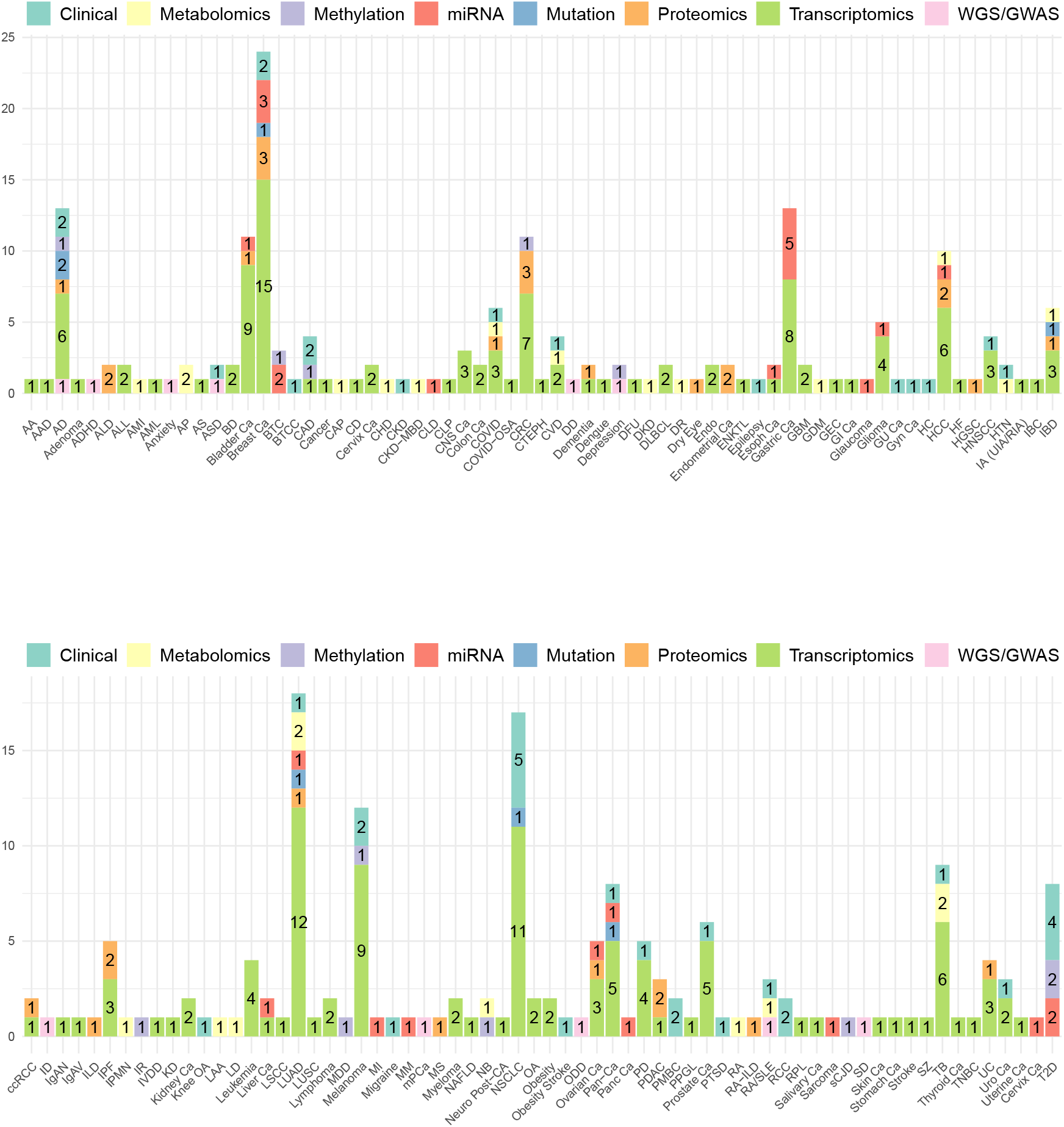
Disease-wise distribution of data modalities.

In contrast, multi-omics approaches have been less frequently used in non-cancer diseases. For instance, COVID-19 research incorporated transcriptomics [146, 147, 196], proteomics [230], metabolomics [196], and clinical data [231], but with fewer studies per modality compared to cancer. Alzheimer’s disease related studies incorporated transcriptomics [138, 139, 232], proteomics [233], methylation [234], and mutation data [203, 204], yet the scope is narrower than in cancers. Moreover, less commonly used modalities, such as WES/WGS/GWAS, are limited to specific diseases, such as autism spectrum disorder [160].

The predominance of multi-omics in cancer research is driven by the disease’s molecular heterogeneity, which involves diverse alterations in gene expression, protein function, and metabolic pathways. Integrating transcriptomics and proteomics data enables the identification of both transcriptional and post-translational biomarkers, as seen in breast cancer [216]. Additionally, the complex tumor microenvironment, involving interactions between cancer and immune cells, necessitates multi-omics to capture comprehensive signatures. Finally, the need for precision medicine in oncology, where biomarkers guide targeted therapies, further underscores the utility of multiomics data as demonstrated in pancancer studies [227]. These factors collectively explain the extensive reliance on multi-omics approaches in cancer biomarker discovery compared to other diseases.

Supplementary Table 3 presents a detailed summary of data splitting strategies used in biomarker discovery studies. Independent testing has remained the most widely used approach, with multiple studies adopting ratios like 80:20 and 70:30. Studies have also frequently used crossvalidation techniques, particularly 10-fold and 5fold CV, to assess model performance. In recent years, many studies have adopted hybrid strategies that combine cross-validation with independent testing to ensure a more rigorous evaluation. Additionally, a few studies have implemented nested CV and leave-one-dataset-out approaches, highlighting efforts to improve model generalizability.

### 5.4 RQ VII: Preprocessing and Feature Engineering Trends Across Modalities and Diseases

This subsection addresses RQ VII by analyzing current trends in preprocessing and feature engineering methods utilized in biomarker discovery studies. We first discuss five distinct categories of preprocessing methods, and then present their distribution across various data modalities. Subsequently, we demonstrate feature engineering categories alongwith their distribution across data modalities and specific diseases. This comprehensive analysis provides researchers with systematic insights into prevalent methodological practices, thereby contributing towards optimal methodological choices and improved biomarker discovery tools.

Figure 9 and supplementary Table 4 illustrate 8 categories of 80 unique preprocessing methods utilized in 69 of 236 studies. Normalization mitigates systematic bias using methods like Quantile normalization [153, 240], Log2 transformation [159, 241], and TMM [242, 243]. Data imputation handles missing values via MICE [244], KNN [215, 245], and MissForest [140]. Batch correction addresses inter-batch variation in data using ComBat [224, 246], MNN, and Harmony [247, 248]. Data cleaning and QC improve data reliability via PCA [249], Shapiro-Wilk tests [232], and duplicate removal [250, 251]. Oversampling tackles class imbalance through SMOTE [164, 252], ADASYN, and ROSE [253, 254]. Background correction reduces technical noise using RMA [142, 255], limma [155], and QC-RFSC [256]. Data alignment and encoding facilitate integration via one-hot encoding [204, 257], ICD simplification [258], and numeric conversion [259]. Lastly, transformation optimizes distributions using Binary Log [260], rank-based inverse normal [161], and CPM normalization [243].

**Fig. 9.**
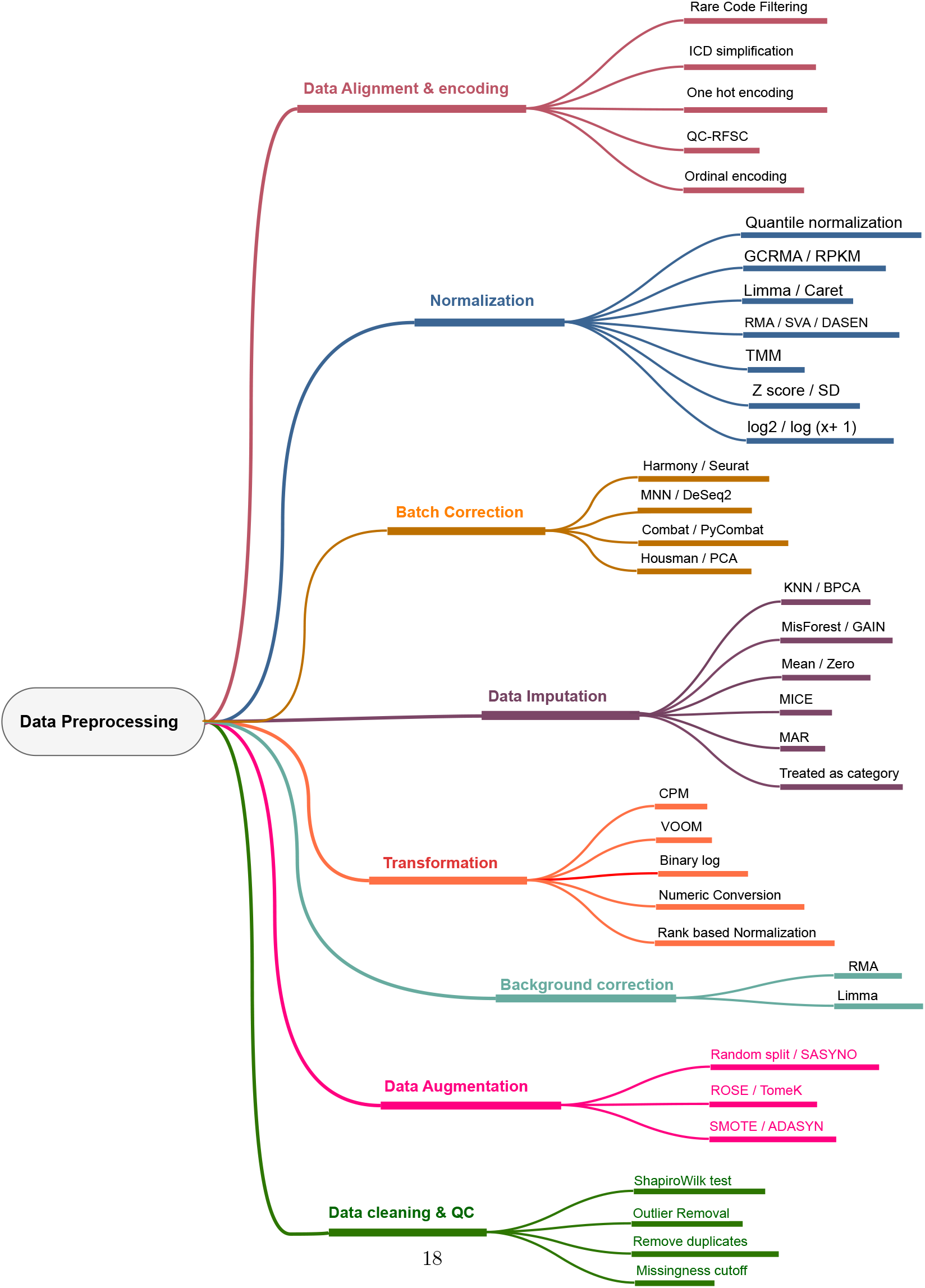
Categorization of preprocessing methods.

Figure 10 and supplementary Table 4 illustrate the utilization of preprocessing methods across 236 biomarker discovery studies. Normalization, data imputation, and batch correction are the most widely used methods due to their critical role in handling bias, missing values, and technical variability. Data cleaning, quality control (QC), and oversampling are also notable. In addition, less frequently applied methods include background correction, data alignment, encoding, and transformation.

**Fig. 10.**
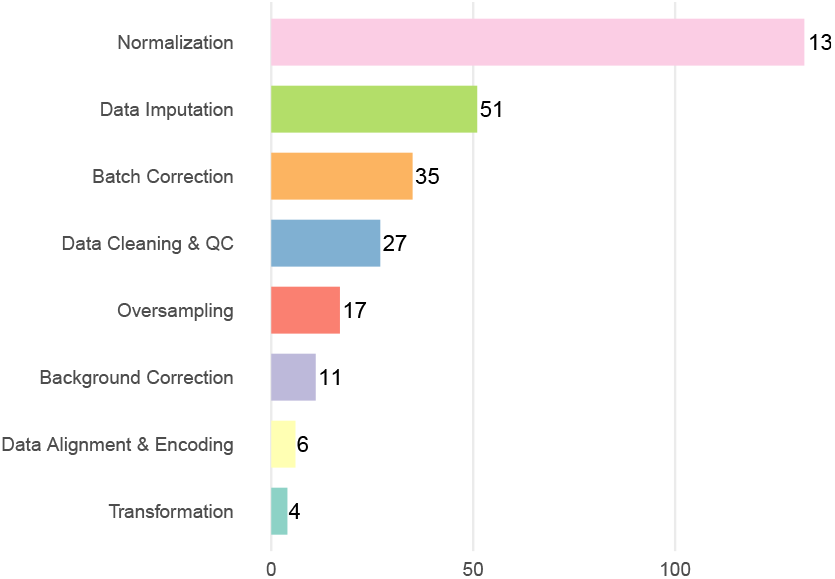
Counts of preprocessing methods used by the reviewed studies.

The selection of the preprocessing methods is tied to the utilized data modalities that are affected by intrinsic noise, sparsity, and batch effects. This highlights the need to investigate which preprocessing methods have been utilized for addressing the modality-specific challenges inherent in diverse omics data types. A thorough analysis of Figure 11 reveals that normalization is the most prevalent preprocessing category across all data modalities, particularly in transcriptomics and methylation data, where standardization is essential to mitigate platform-induced biases and ensure comparability across samples. Data imputation follows closely and is prominently adopted in clinical, transcriptomic, and metabolomic studies to fill missing values. Batch correction is applied to transcriptomics and methylation data, due to their susceptibility to technical variability across platforms and batches. Data cleaning and quality control methods are moderately applied across modalities, with greater emphasis in clinical and proteomics data. In contrast, background correction, alignment, and encoding are less frequent, primarily observed in transcriptomics and miRNA studies. The minimal use of these methods in WGS and mutation data suggests either alternative preprocessing pipelines or a lack of consensus in best practices. Overall, the heterogeneous distribution of preprocessing methods underscores the necessity of tailoring strategies to data-specific challenges to enhance robustness and performance of downstream biomarker discovery tools/pipelines.

**Fig. 11.**
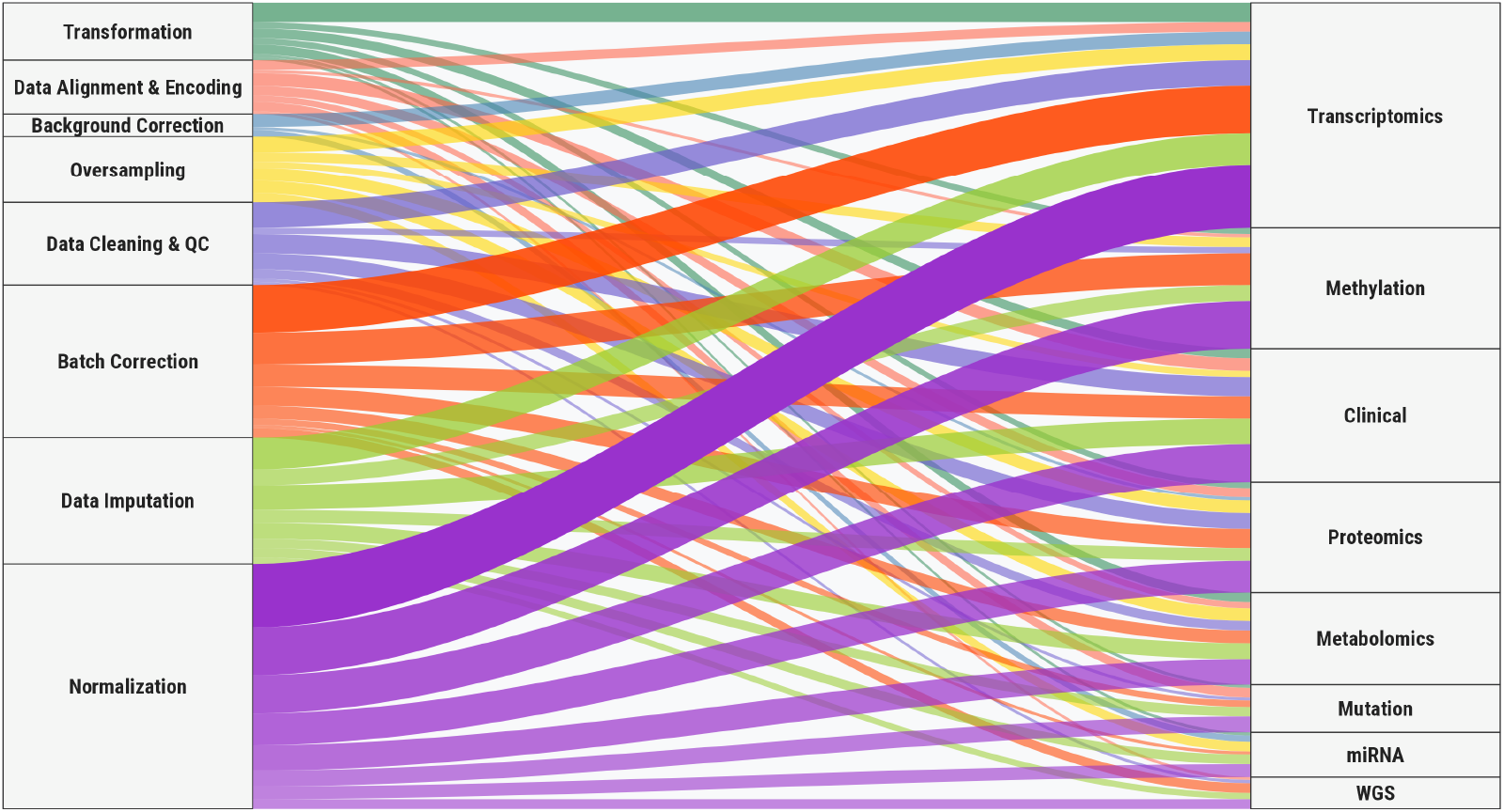
Utilization of preprocessing methods in biomarker discovery studies across 8 distinct data modalities.

Figure 13 illustrates the top 35 feature engineering methods utilized in biomarker discovery studies, which can be broadly categorized into two primary paradigms: dimensionality reduction and feature selection. Dimensionality reduction methods aim to transform high-dimensional omics or clinical data into a more compact and informative representation and can be further grouped into linear, nonlinear, and matrix factorization-based methos. Linear methods include PCA [134, 261], Independent Component Analysis (ICA) [211], sparse PCA, and Partial Least Squares /Partial Least Squares Discriminant Analysis (PLS/PLSDA) [149, 240, 262–264]. Nonlinear methods comprise Autoencoders [131, 202, 252], Variational Autoencoders (VAEs), Uniform Manifold Approximation and Projection (UMAP) [265, 266], and random projection [267]. Additionally, matrix factorization-based methods include Non-negative Matrix Factorization (NMF) [268] and negative binomial models [242].

Feature selection methods aim to identify a subset of informative features from the original data without applying transformation and can be systematically categorized into several subgroups: statistical, information-theoretic, interpretability-based, wrapper, embedded, graph- and network-based, expert-driven, correlation-based, heuristic and metaheuristic methods. Statistical methods include t-tests [125, 200, 269, 270], Analysis of Variance (ANOVA) [163, 236, 271], Wilcoxon rank-sum test [125, 163], Kruskal–Wallis test [227], Mann–Whitney U test [141], chi-square test [203], Fisher’s exact test [272, 273], differential expression analysis using DESeq2 [163], edgeR [243], and fold-change [269]. Information-theoretic and interpretability-based methods comprise mutual information [203, 274], Joint Mutual Information (JMI) [166], SHAP [275], DL Important FeaTures (DeepLIFT) [252], TreeSHAP [276], Integrated Gradients [252], graph-based ranking, and SelectKBest [128]. Wrapper and embedded approaches include Least Absolute Shrinkage and Selection Operator (LASSO) [127, 130, 145, 146, 156, 221, 277, 278], elastic net [143, 237], ridge regression [143, 237], RFE [153, 154, 199, 271, 279], Boruta [147, 280, 281], Generalized Linear Models with Stepwise Akaike Information Criterion (GLMStepAIC) [282], Tree-based Pipeline Optimization Tool (TPOT) [283], Hilbert-Schmidt Independence Criterion Lasso (HSIC Lasso) [284], Variable Importance in Projection for PLS-DA (VIP-PLSDA) [256], Smoothly Clipped Absolute Deviation (SCAD) [285], and regularized generalized linear models [121]. Graph- and network-based strategies include Weighted Gene Co-expression Network Analysis (WGCNA) [156, 278, 278], graph-based centrality measures [190], Multiscale Embedded Gene Co-expression Network Analysis (MEGENA) [277], intramodular connectivity [286], and Gene Regulatory Networks (GRNs) [120]. Biologically informed or expert-driven techniques comprise GeneMANIA [287], PLINK [288], Minor Allele Frequency (MAF) thresholds [195], literature-guided selection [**? ?**], Orchid’s feature selection [289], and expert- or data-driven curation [213]. Correlation-based methods include Pearson correlation [259], Spearman rank correlation [161], and cosine similarity [272]. Heuristic and metaheuristic algorithms involve approaches such as the Grey Wolf algorithm [290], Genetic Algorithms [137], Random Drift Optimization [291], Sequential Forward Selection [292], and bootstrapping [209]. Finally, clustering-based feature engineering methods include hierarchical clustering [152], K-means clustering [235], and Iterative Local Refinement Clustering (ILRC) [158].

Figure 12 presents the distribution of the top feature engineering methods utilized in biomarker discovery studies. It highlights more frequent use of methods like LASSO, feature importance, and RFE. Classical statistical tests such as t-test, ANOVA, and PCA also remain prevalent, which reflects their foundational role in biomarker discovery studies. Emerging approaches from DL and bioinformatics—such as autoencoders, WGCNA, and ssGSEA—illustrate growing diversity in methodological choices.

**Fig. 12.**
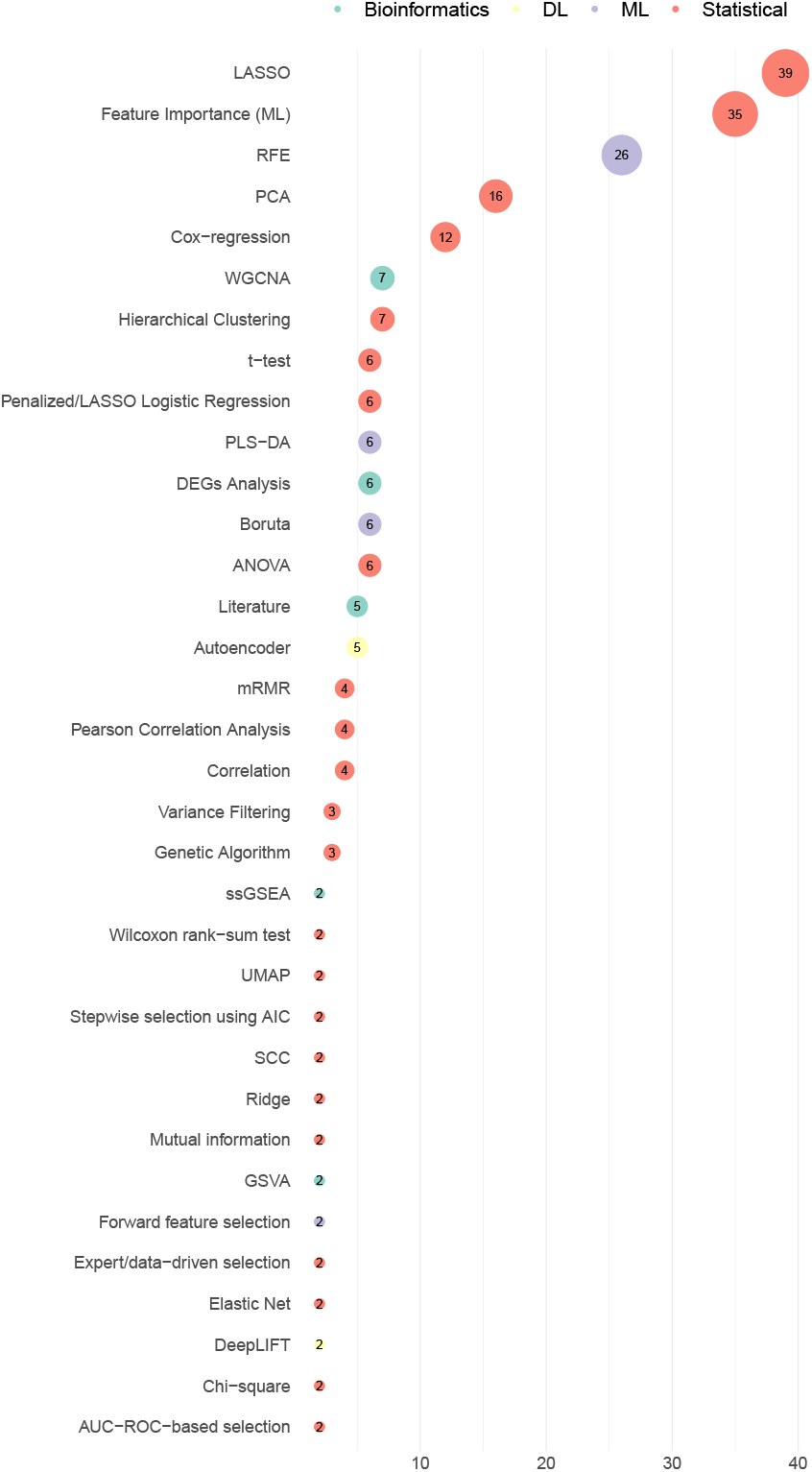
Counts of feature engineering methods used by the reviewed studies.

**Fig. 13.**
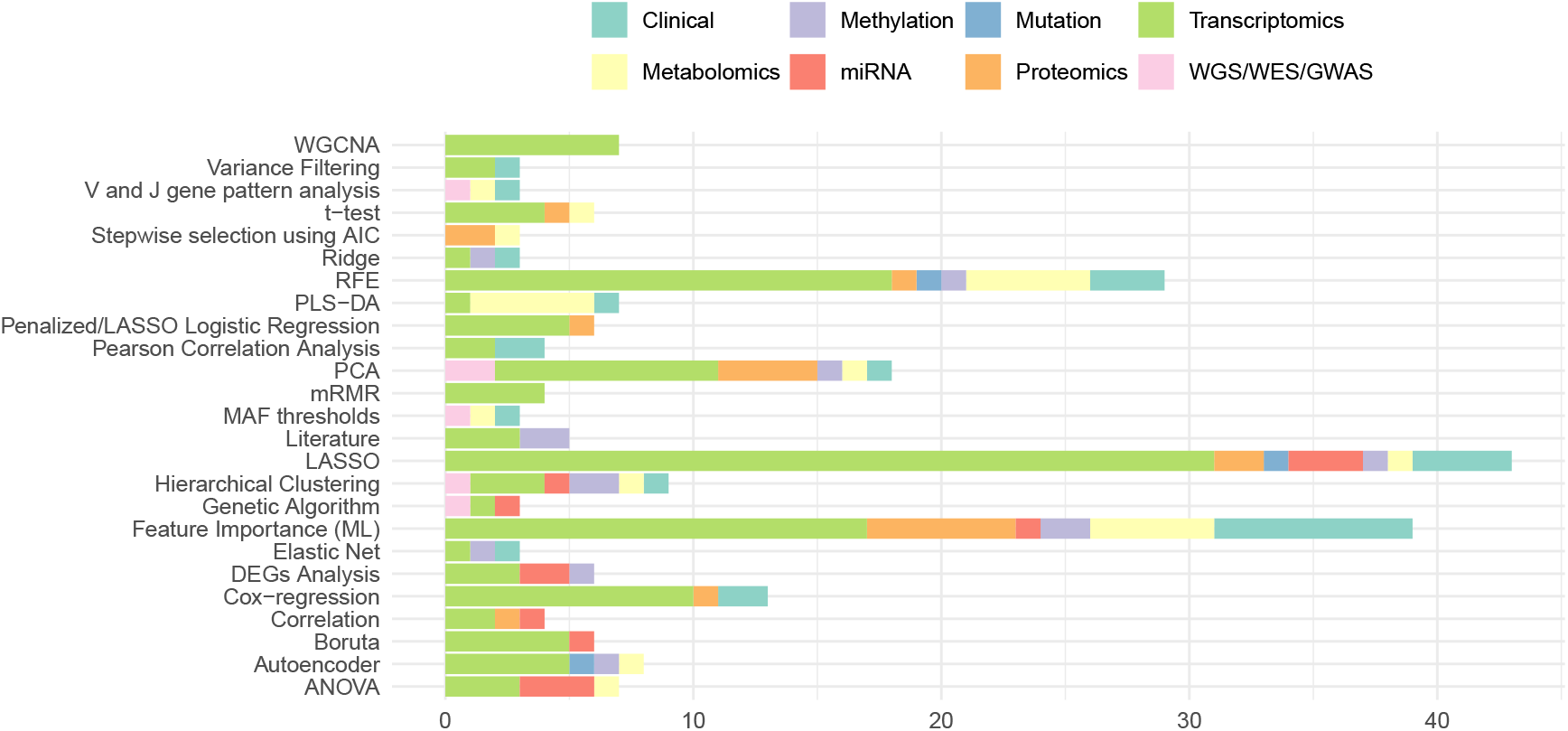
Distributiuon of feature engineering methods across 8 distinct data modalities.

An in-depth analysis of feature engineering methods used in biomarker discovery reveals distinct patterns in their applicability across various data modalities, including transcriptomics, proteomics, methylation, metabolomics, mutation, miRNA, clinical, and WGS/WES. Among all methods, LASSO, FI, RFE, and PCA stand out as the most broadly applied across all data modalities. These methods offer flexibility in handling both structured and high-dimensional data, which explains their widespread adoption. Conversely, several methods are modality specific, i.e., DESeq2, edgeR, and fold-change analysis have only been applied to transcriptomic data. Likewise, GSVA, ssGSEA, and RNs methods are used exclusively in omics data modalities. DL approaches like DeepLIFT, TreeSHAP, and AE have seen growing interest but remain largely limited to transcriptomics and methylation, with limited adoption in proteomics and clinical datasets. Some methods, despite their theoretical potential, remain unexplored or underutilized in specific modalities. For instance, SHAP, IG, and Graphbased ranking methods have not been applied to methylation, proteomics, or miRNA datasets in the reviewed studies. Furthermore, advanced correlation-based techniques such as MIC, SPCC, KCC, and PPCC are found only in transcriptomics.

### 5.5 RQ VIII: Trends of ML/DL Models Across Diverse Diseases, and Modalities

In pursuit of addressing RQ VIII, this section presents an overview and insights into ML and DL models utilized in biomarker discovery studies. It examines emerging methodological trends across a diverse spectrum of diseases and aims to support researchers in identifying underexplored areas and enhancing predictive modeling strategies.

Figure 14 illustrates the distribution of ML and DL models across 236 diverse biomarker discovery studies. Traditional ML algorithms, particularly RF, SVM, logistic regression (LR), LASSO, and DT, have been widely used. Gradient-boosted models such as XGBoost (XGB) and GB also show notable usage. In contrast, DL methods, including convolutional neural networks (CNNs), and deep neural networks (DNN) exhibit moderate adoption, primarily limited by data availability and complexity constraints. Survival-oriented models such as Cox Proportional Hazards (CoxPH), CoxBoost, and Random Survival Forests (RSF) maintain specialized roles for time-to-event predictive analyses.

**Fig. 14.**
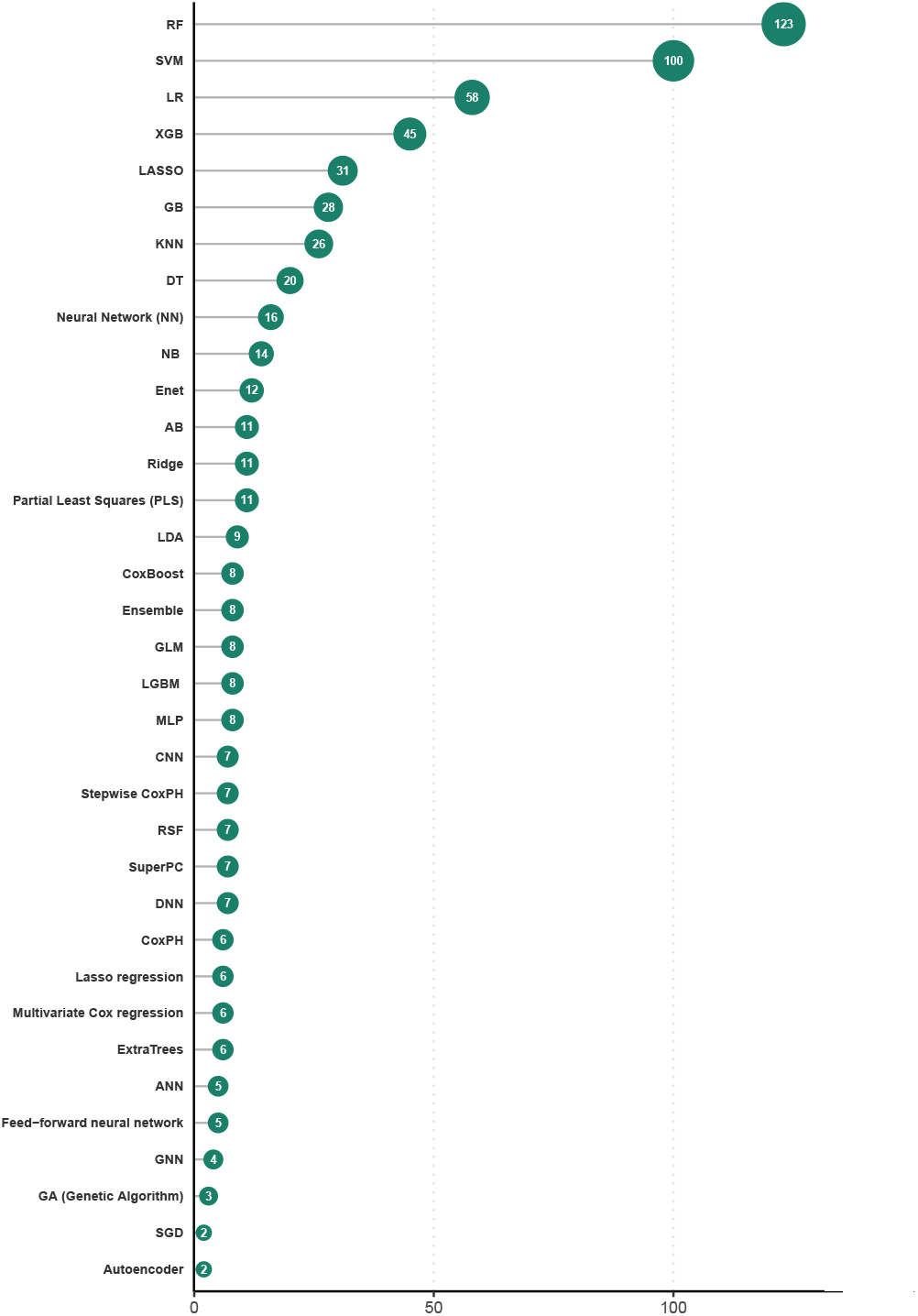
Distribution of ML and DL models across biomarker discovery studies.

Supplementary Table 5 presents the utilization of 58 unique ML and DL models across 142 distinct diseases. In terms of ML, SVM has been frequently used in hepatocellular carcinoma, colorectal cancer, breast cancer, and bladder cancer studies [123, 141, 147, 150, 191, 210, 211, 216, 279, 293, 294]. RF has been commonly used for colorectal cancer, lung cancer, and COVID- 19 [150, 162, 210], while LR has appeared in studies on breast cancer, Kawasaki disease, and lung disorders [123, 150, 162]. In recent years, DL models have gained traction, although their usage remains relatively limited. Artificial Neural Networks (ANN), CNNs, and hybrid architectures such as CNN-RNN and CNN-LSTM have been explored in diseases like lung cancer, glioblastoma multiforme (GBM), Alzheimer’s disease, and head and neck squamous cell carcinoma (HNSC) [145, 159, 239, 252, 295].

Figure 15 presents the distribution of ML and DL models across eight distinct data modalities. A comprehensive analysis reveals that transcriptomics and clinical modalities have been extensively explored, utilizing 40 and 29 unique ML and DL methods, respectively. In contrast, other modalities, such as mutation (14 models), miRNA (15 models), and WGS/WES/GWAS (15 models), exhibit limited methodological diversity, which indicates insufficient exploration and highlights the potential for future research efforts. Notably, LR, SVM, neural networks (NN), and RF have been adopted across all modalities ([123, 155, 296]). Conversely, several specialized methods, such as Gaussian Bayes, Quadratic Discriminant Analysis (QDA), CapsuleNet, and SuperPC, remain modality-specific, having been used within only a single data modality [224, 296, 297]. These trends underscore critical gaps and opportunities for extending the application of diverse ML and DL methods across less explored data modalities.

**Fig. 15.**
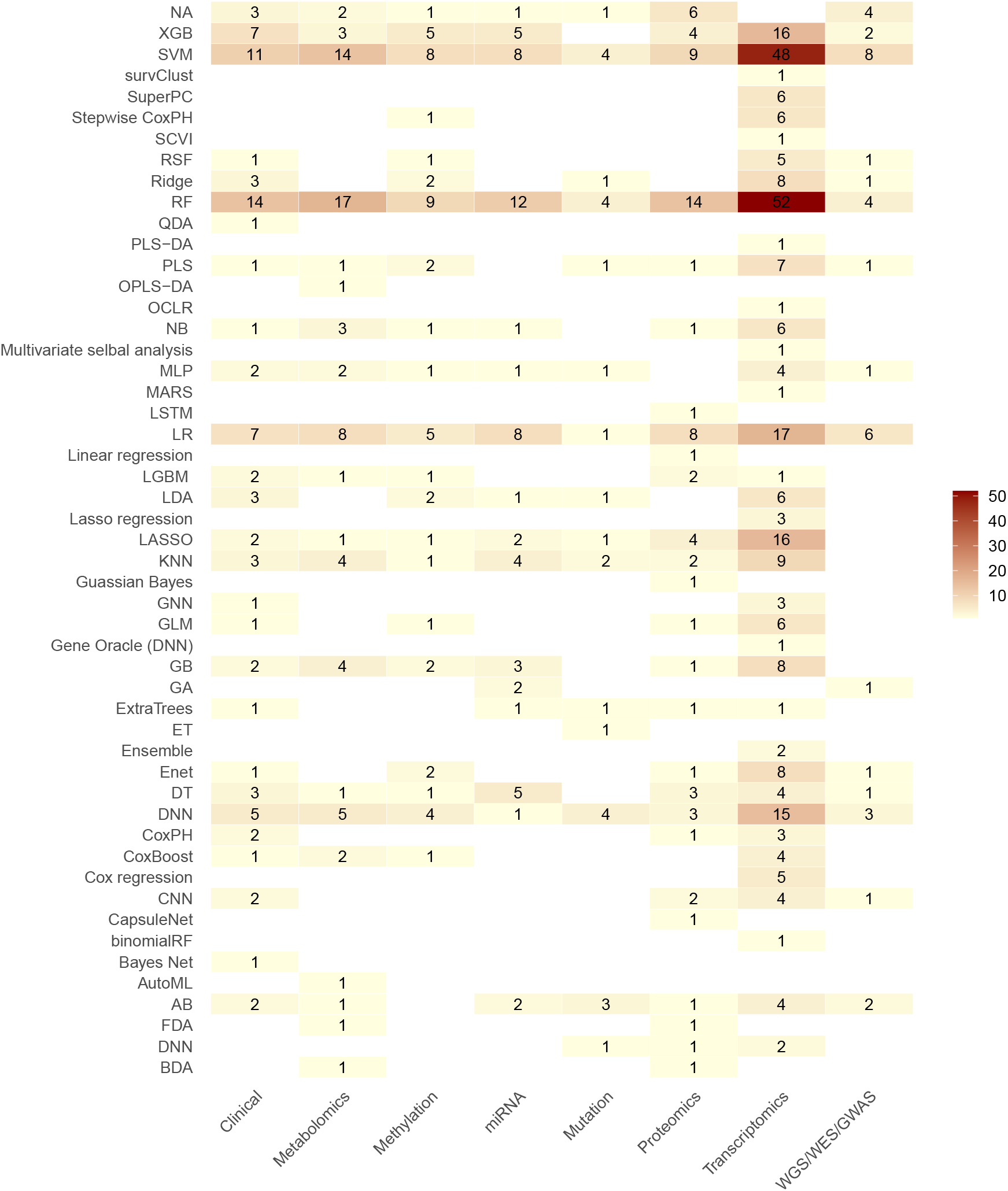
Distribution of ML and Models in terms of 8 distinct omics modalities.

### 5.6 RQ IX: Open Source Biomarker Discovery Studies

Following RQ IX, this section provides a detailed summary of publicly accessible source codes developed for biomarker discovery. This comprehensive overview enables researchers to advance existing work, promoting collaboration and the rapid development of biomarker discovery tools.

Supplementary Table 5 presents an extensive compilation of open-source 88 studies that utilized a range of ML, DL, and statistical methods. A thorough investigation reveals that only 66 of these studies made their codebase publicly available, while the remaining 24 studies stated code availability upon request.

Among the publicly shared resources, Python is the most preferred development environment, with over 70% of tools built using TensorFlow, PyTorch, Scikit-learn, XGBoost, or Keras frameworks [128, 148, 150, 220, 252, 298]. On the other hand, a smaller but notable proportion of studies employed R, leveraging packages such as caret, glmnet, ranger, and survival for classical ML and survival modeling tasks [189, 241, 299].

A thorough analysis reveals that only a minority of these tools were built using standardized biomarker discovery frameworks. Instead, many studies opted to develop custom workflows or hybrid ensembles, combining multiple classifiers (e.g., RF, SVM, XGBoost, NN) and integrating dimensionality reduction, feature selection, and hyperparameter tuning [141, 147, 148]. Notably, a few models incorporated explainability modules using gradient-based or SHAP-style interpretation to identify high-confidence biomarkers or decision-driving features by using SHAP or Captum libraries [244, 252, 300].

While these open-source tools present an invaluable foundation for reproducible science and collaborative advancement, the lack of standardized code sharing protocols, inconsistent documentation, and missing specifications remain critical bottlenecks. Researchers aiming to reuse or extend these pipelines should consider benchmarking across curated datasets and clearly defined evaluation metrics. The availability of these open-source libraries, as outlined in supplementary Table 5, not only fosters transparency and reusability but also accelerates innovation in AI-driven biomarker discovery.

### 5.7 RQ (X): Clinical and Translation Potential of Predicted Biomarkers

This subsection addresses RQ X through a structured categorization of diseases and associated biomarkers across 19 distinct therapeutic areas. It further examines the distribution of diseases and biomarkers within these categories to identify therapeutic areas with extensive exploration.

Figure 16 presents the categorization of 147 diseases along with counts of biomarkers in 19 different therapeutic areas. Oncology emerges as the most explored therapeutic area, encompassing 60 diseases with 1,163 unique biomarkers across numerous studies. These include extensively studied cancers such as breast cancer [118, 120, 227], non-small cell lung cancer (NSCLC) [201, 239], and hepatocellular carcinoma [140]. Following oncology, neurology, and cardiology have also been well explored, with 14 and 13 diseases, respectively, with a major focus on Alzheimer’s and Parkinson’s diseases [138, 232, 251]. In cardiology, diseases such as acute myocardial infarction [144], coronary artery disease [288], and aortic dissection [145] have been explored. Overall, in cardiology and neurology, around 256 biomarkers have been identified with AI-driven biomarker discovery. Psychiatry, rheumatology, and infectious diseases follow with 11, 7, and 6 diseases with a major focus on schizophrenia, bipolar disorder [139, 141, 210], COVID-19, and Tuberculosis [146, 147, 301]. On the other hand, several therapeutic domains remain underrepresented. Areas such as orthopedics, gynecology, ophthalmology, neurosurgery, and Dermatology are each associated with only 1–2 diseases and a limited number of discovered biomarkers. Despite significant advancements in AI-based biomarker discovery, there is an urgent need for systematic, large-scale wet lab validation pipelines that can experimentally assess the reliability, biological relevance, and translational potential of these AI-predicted biomarkers in real-world clinical settings.

**Fig. 16.**
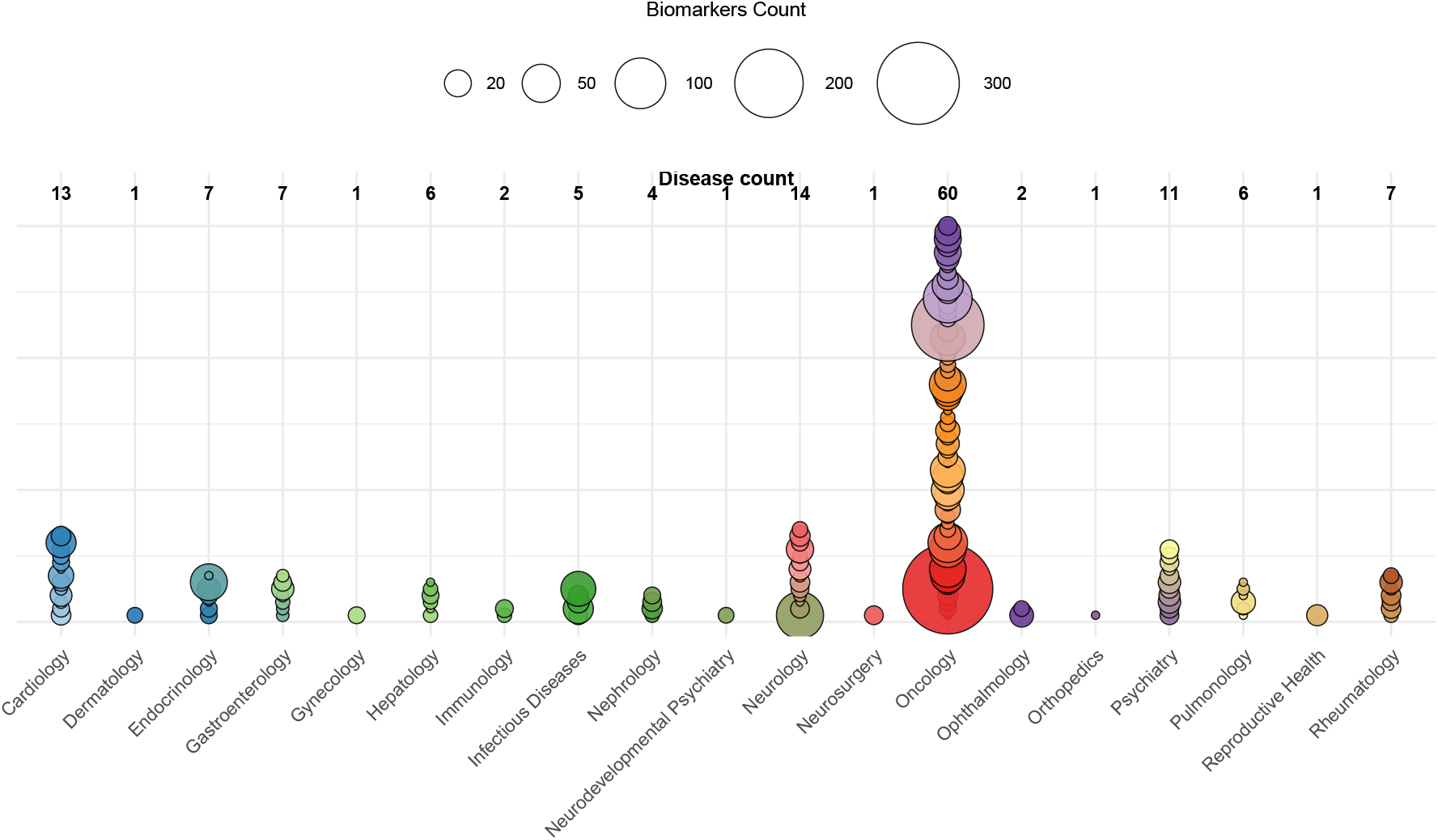
Categorization of diseases and biomarkers in 19 distinct therapeutic areas.

### 5.8 RQ XI: Publisher and Journal-Wise Distribution of Research Papers

To address Research Question XI, Figures 17 and 18 present the distribution of biomarker discovery studies across a range of journals and publishers. This analysis helps researchers identify strategic publication venues and encourages interdisciplinary collaboration in the field of biomarker discovery. Approximately one-third of the studies have been published through Springer. Within Springer, a variety of domain-specific journals such as BMC Cancer, BMC Genomics, BMC Bioinformatics, and BMC Medical Genomics appear prominently. Nature Publishing Group follows with around 70 publications, through journals such as Nature Communications and NPJ Digital Medicine. Elsevier has published around 20 relevant studies through journals such as Computers in Biology and Medicine and Clinical Epigenetics. Other publishers include Frontiers Media, MDPI, Wiley Online Library, and Oxford University Press, each contributing between 5 and 20 studies. In total, the studies are distributed across 25 different publishers and over 80 distinct journals.

**Fig. 17.**
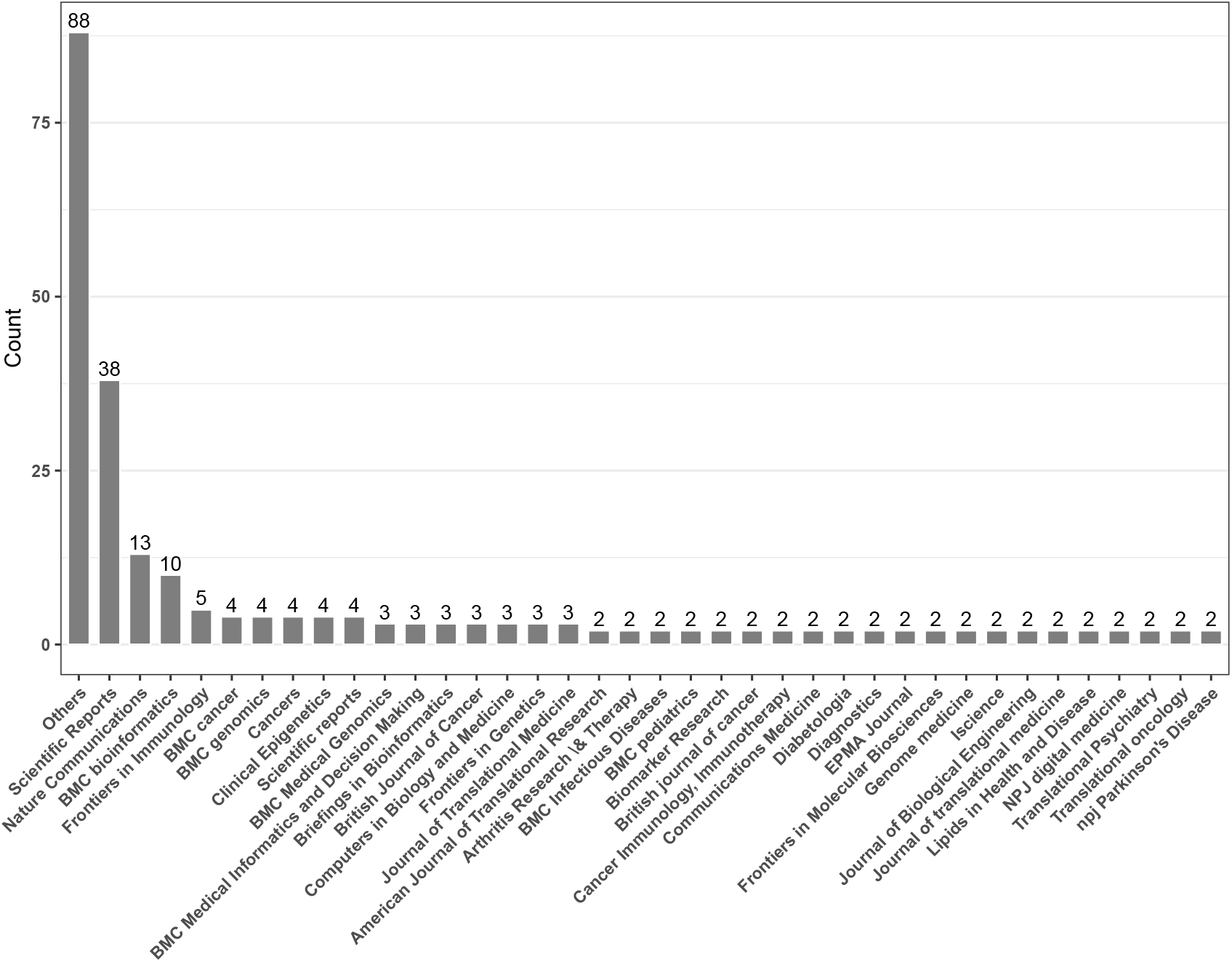
Journal-wise distribution of research articles.

**Fig. 18.**
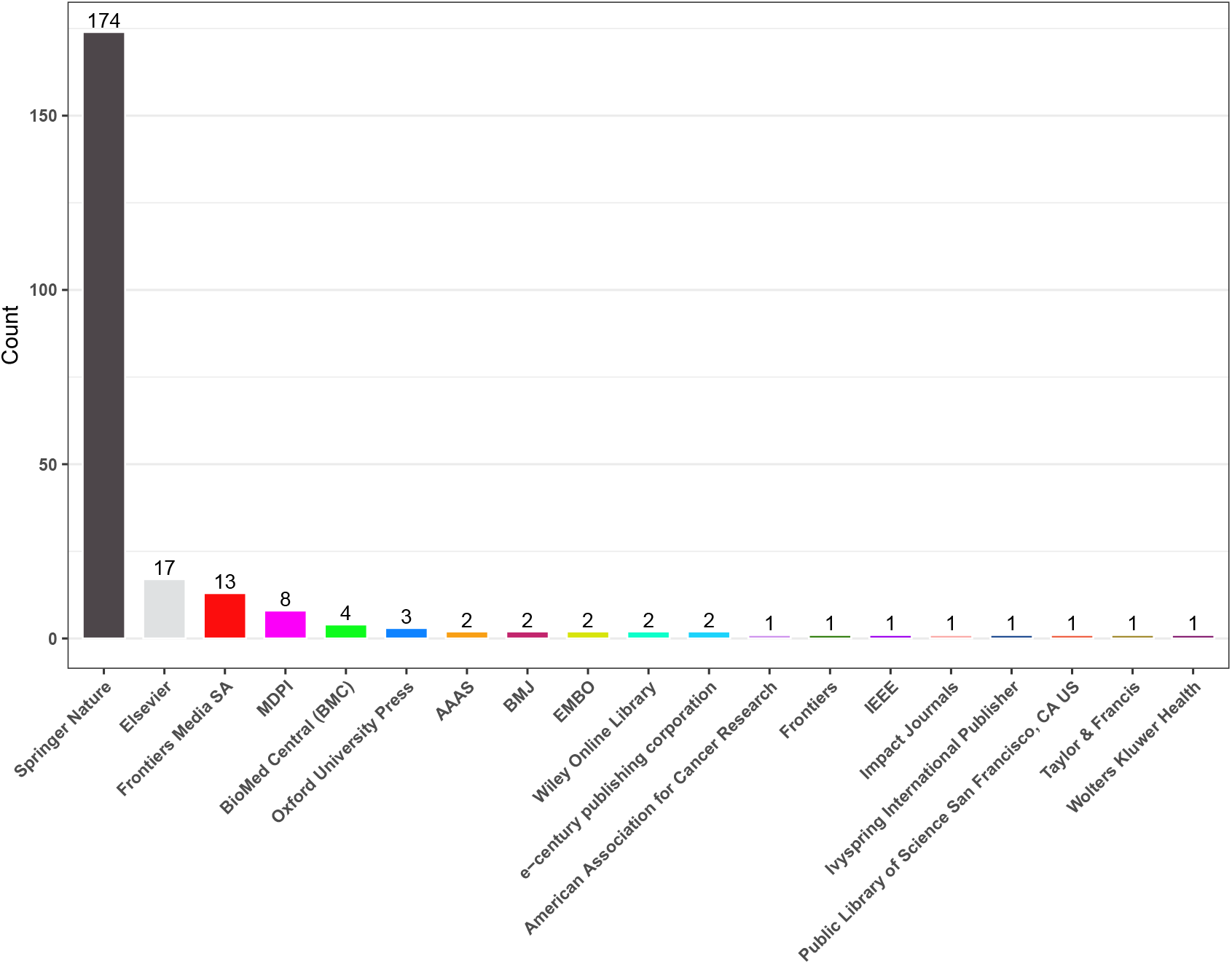
Publisher-wise distribution of research articles.

## 6 Discussion

The field of AI-driven biomarker discovery has emerged as a cornerstone in advancing precision medicine by enabling early diagnosis, disease stratification, and treatment monitoring. With the increasing number of complex diseases and the need for individualized care, there is a growing demand and development of disease specific AI-driven biomarker discovery tools. Our systematic review unveils that researchers put dominant research focus on cancer, with breast [118–121, 302], lung [129, 163, 201, 221, 239, 266, 274, 303], gastric [121, 125, 126, 208], and skin cancers [128, 129, 266, 296, 303] being among the most extensively studied. This concentration is rational, given the molecular heterogeneity of cancers and the critical need for subtype-specific biomarkers in clinical oncology. Additionally, the increasing availability of multi-omics cancer datasets from public resources like TCGA [54] and GEO [55] has accelerated cancer-centric research.

While oncology dominates the biomarker discovery landscape, researchers are extending it into other therapeutic domains such as neurology, cardiology, infectious diseases, and autoimmune disorders. However, these areas are relatively underexplored, often constrained by limited availability of high-quality, multimodal datasets. For instance, despite their substantial global impact, cardiovascular [161, 202, 271] and neurodegenerative diseases [138, 212, 268, 304] remain underrepresented in AI-driven biomarker discovery research due to the limited availability of high-quality datasets. This limitation presents opportunities for targeted efforts to expand biomarker discovery into non-cancer diseases, especially by promoting dataset standardization and accessibility.

Access to well-annotated public and private datasets serves as a foundational pillar for advancing AI-driven biomarker discovery. Our analysis reveals that the majority of studies rely on public repositories such as GEO, TCGA, ArrayExpress, and ENCODE, which provide rich, multi-omics datasets across a wide spectrum of diseases, particularly in oncology. These datasets facilitate model training, reproducibility, and benchmarking. However, a significant number of studies also utilize private or in-house datasets—especially for rare diseases, clinical trials, or proprietary diagnostic platforms—highlighting the need for secure data-sharing frameworks. Despite their value, private datasets often lack standardized formats and accessibility, which limits their broader validation and reuse. Bridging the gap between public and private data ecosystems through federated learning and harmonized metadata standards could significantly advance the field by enabling more robust and generalizable biomarker discovery pipelines.

One of the defining features of biomarker discovery is the integration of diverse data modalities, including transcriptomics, methylation, proteomics, and clinical data. Our findings reveal a strong reliance on transcriptomics, proteomics, and clinical data, particularly in cancer research, while data types such as GWAS, WGS, and WES remain underutilized. Moreover, multimodal integration remains a challenge, with only a handful of studies combining 3 or more data modalities. Effective multimodal data integration is essential for capturing the full biological complexity of disease mechanisms and holds promise for discovering more comprehensive and actionable biomarkers.

A key insight from our analysis is the centrality of preprocessing methods in ensuring the reliability of downstream analysis. Methods like normalization [192, 198, 220, 224, 234, 305], data imputation [196, 213, 244], and batch correction [118, 136, 236] were frequently used to address technical variability and missing values. Notably, the choice of preprocessing methods often varies by modality—for instance, batch correction is predominantly applied in transcriptomics and methylation datasets, while imputation is more common in clinical and metabolomic data. However, many studies overlook rigorous quality control procedures or fail to report them transparently, potentially undermining reproducibility and generalizability. Therefore, standardized and well-documented preprocessing pipelines are essential for enhancing model robustness.

Feature engineering plays a vital role in biomarker discovery, with a wide array of dimensionality reduction and feature selection methods being used. Our analysis shows that traditional statistical methods such as PCA, t-tests, and ANOVA remain widespread, while modern methods like autoencoders and perturbation/gradient-based explainability are gaining traction. Nonetheless, few studies benchmark multiple feature engineering methods under the same settings, making it difficult to conclude the optimal strategies. Large-scale benchmarking efforts would thus provide valuable guidance for the selection of appropriate methods tailored to specific data types and diseases.

In terms of ML and DL models, our review identifies the widespread use of classical ML algorithms namely, SVM [150, 191, 210, 221, 279, 294], RF [127, 137, 164, 208, 235, 236, 299, 306], LR [120, 123, 125, 221, 294] alongside an increasing adoption of NN-based models [122, 141, 145, 148, 189**?**]. While DL models offer superior performance in handling complex non-linear interactions, they demand large datasets and remain limited by their “black-box” nature. In contrast, ML models are more interpretable and work effectively on smaller datasets. Striking a balance between model complexity and interpretability is crucial, especially in clinical contexts where explainability is vital for trust and adoption.

XAI has emerged as a critical component in the biomarker discovery pipeline. Although many studies recognize the importance of explainability, limited studies incorporate XAI techniques such as SHAP [196, 275], LIME, or DeepLIFT [307], and integrated gradients [252, 307]. The application of XAI helps unravel the inner workings of ML/DL models, offering insights into the biological relevance of selected features and facilitating clinical translation. However, most studies implement post hoc explainability techniques rather than developing inherently explainable models. We advocate for the broader adoption and benchmarking of XAI methods, particularly in studies intended for clinical deployment.

Despite the availability of XAI tools, current biomarker discovery studies do not evaluate the reliability of feature attributions derived from black-box models. Techniques like SHAP or Integrated Gradients often highlight important genes or molecular signatures, yet none of the studies assess whether these attributions are consistent, replicable, or biologically meaningful. Attribution reliability remains a critical missing link in establishing trust in AI-discovered biomarkers. Hereby, we advocate for the incorporation of systematic XAI evaluation measures to rigorously assess the reliability and utility of model attributions in biomarker discovery. Metrics such as Area Over the Perturbation Curve (AOPC) [308, 309] provide insight into how effectively the identified features contribute to model predictions by quantifying performance drop when top features are masked. Continuity [308, 309] evaluates whether small perturbations in input lead to stable explanations, thereby testing explanation robustness. Infidelity [308, 309] measures the degree to which an attribution method captures the true causal effect of input features on the model output, offering a critical test of faithfulness. By integrating these evaluation strategies, researchers can move beyond superficial interpretation and begin to establish confidence in the biological relevance, reproducibility, and trustworthiness of AI-derived biomarkers.

Evaluation of model performance is another critical aspect that lacks standardization. Common metrics such as accuracy, and precision are extensively used, but often fail to capture nuances relevant to imbalanced datasets or survival outcomes. Advanced metrics like integrated Brier score [310], time-dependent AUC, or calibration measures are rarely used but could offer more robust evaluations, especially in longitudinal or survival analyses. Utilizing a suite of complementary metrics can ensure more comprehensive model assessment and better support real-world applications.

Despite the growing use of AI in biomarker discovery, the majority of studies have not shared their code or analysis pipelines. This lack of code availability hampers transparency, reproducibility, and independent validation—core principles of scientific rigor. Without access to the underlying implementation, it becomes challenging to assess the reliability of reported results, replicate findings across cohorts, or adapt methods to new datasets. Consequently, the absence of opensource code limits collaborative progress and slows the translation of computational biomarkers into clinical and experimental applications.

Finally, the translational potential of AI-discovered biomarkers remains an open challenge. Despite numerous discoveries reported in the literature, few biomarkers have advanced to clinical validation or regulatory approval. Barriers include limited external validation, small cohort sizes, lack of reproducibility, and insufficient integration with clinical workflows. Addressing these gaps will require multi-institutional collaborations, public data sharing, and rigorous validation protocols, including prospective and multicentric studies. Furthermore, harmonizing biomarker discovery pipelines with clinical standards will enhance their utility in diagnostics, prognostics, and treatment personalization.

In conclusion, AI-driven biomarker discovery represents a transformative frontier in precision medicine. Our review reveals both the breadth of current research and the critical challenges that remain. Future studies must emphasize multimodal integration, preprocessing standardization, benchmarking of feature engineering strategies, inclusion of XAI, and rigorous model validation to bridge the gap between computational discoveries and clinical impact.

